# Changes in microglia chromatin accessibility in aged female mice

**DOI:** 10.1101/2024.01.18.575944

**Authors:** Victor A. Ansere, Kyla B. Tooley, Kottapalli Srividya, Walker Hoolehan, Kevin D. Pham, Sarah R. Ocañas, Willard M. Freeman

## Abstract

Aged female microglia display a more inflammatory and disease-associated phenotype compared to age-matched males. Epigenetic mechanisms, such as chromatin accessibility, are key drivers of microglial plasticity and phenotypes necessary for development, priming, and immune activation. Therefore, alterations in chromatin accessibility patterns can potentially regulate the neuroimmune responses and phenotypes observed in female microglia with aging, but to date have not been assessed. In this study, hippocampal microglia chromatin accessibility in young (4-5 months) and old (23-24 months) female mice was interrogated by Assay for Transposable Accessible Chromatin using Sequencing (ATAC-Seq). Cx3cr1-cre/ERT2+: NuTRAP mice were used to tag microglia and enable INTACT (isolation of nuclei tagged in specific cell types) collection of microglia-specific nuclei. With aging, loci specific gains and losses in chromatin accessibility were observed. Notably, changes in chromatin accessibility were skewed, with aged female microglia having more regions gaining accessibility than loosing accessibility. These changes were under-represented in the proximal promoter region (≤1kb) of genes but were enriched in intergenic regions. Regions that gained accessibility were more concentrated around genes responsible for myeloid cell differentiation and the regulation of immune and inflammatory responses. In contrast, regions that became less accessible were closest to genes involved in neuronal and synaptic function. In addition, X Chromosome accessibility changes were less common compared to autosomal changes, which argues against increased X Chromosome escape from inactivation with aging in female microglia. Overall, our data demonstrate age-related chromatin accessibility changes in female microglia, which may be regulated within enhancers and distal regulatory elements, and that these changes have potential downstream implications for the inflammatory phenotype of microglia in aging female mice.

## 1. Introduction

Microglia serve as the primary resident immune cells of the brain and are involved in maintaining brain homeostasis through the initiation of immune responses and the removal of neuronal debris ^1,2^. Initially, microglia were thought to transition between only two morphological states, ramified to amoeboid, which determined their functional or reactivational phenotypes ^3^. However, recent advances indicate that distinct spectrums of gene expression profiles occur throughout the lifespan of microglia including aging, whether in a ‘steady’ state or in response to stimuli ^4–6^. It has been hypothesized that microglia populations have a slow self-renewal rate ^7–9^, making them susceptible to accumulating age-related effects. Age-related transcriptomic analysis within the hippocampus reveals cell-type specific signatures, with microglia-specific genes dominating age-related changes ^10^. Notably, these aging effects are more pronounced in females ^5,10–12^. The impact of aging on microglia translates into impaired phagocytic activity, neuroprotective functions, and an elevated expression of proinflammatory cytokines, which collectively characterize a state of chronic inflammation in the brain ^7,13^.

Epigenetic patterns, which include DNA and histone modifications as well as chromatin accessibility, are emerging as important mechanisms that modulate genome accessibility and transcription programs necessary for the diverse range of microglia signatures ^14–16^. These mechanisms have been suggested to regulate microglia polarization towards specific phenotypes, as well as the long-lasting effects of stimuli essential for processes like microglial priming and maternal immune activation ^14–17^. Together with changes in the patterns of DNA modifications and chromatin marks, altered chromatin accessibility is a hallmark of the aging process ^18,19^, potentially contributing in a cell type-specific fashion to the age-related structural and functional decline of brain function. Although the potential role of age-related epigenetic changes on microglial function is acknowledged, studies examining alterations in microglial chromatin accessibility with aging are lacking.

In a previous aging transcriptomic study, we observe that female microglia are more skewed to disease-associated and senescent phenotypes, even in the absence of disease pathology ^12^. Thus, studying chromatin accessibility in female microglia will provide a more thorough understanding of how epigenetic programs are disrupted with aging and disease. We isolated microglia nuclei from both young (4-5 months) and old (23-24 months) female Cx3cr1-cre/ERT2+: NuTRAP ^20^ (The Nuclear Tagging and Translating Ribosome Affinity Purification) mice through an Isolation of Nuclei TAgged in specific Cell Types (INTACT) approach **(Figure 1A & B)**. Using the Assay for Transposase-Accessible Chromatin using sequencing (ATAC-seq) ^21,22^ **(Figure 1C)**, we identified open chromatin regions and compared the differences between the two age groups. We sought to address the following questions: 1) What changes are occurring to the chromatin accessibility landscape in aging female microglia; 2) how are these changes distributed across various genomic locations; 3) how do chromatin accessibility changes relate to alterations in the transcriptomic profile of female microglia; and 4) how do changes in chromatin accessibility on the X chromosome compare with those on autosomes?

**Figure 1.**
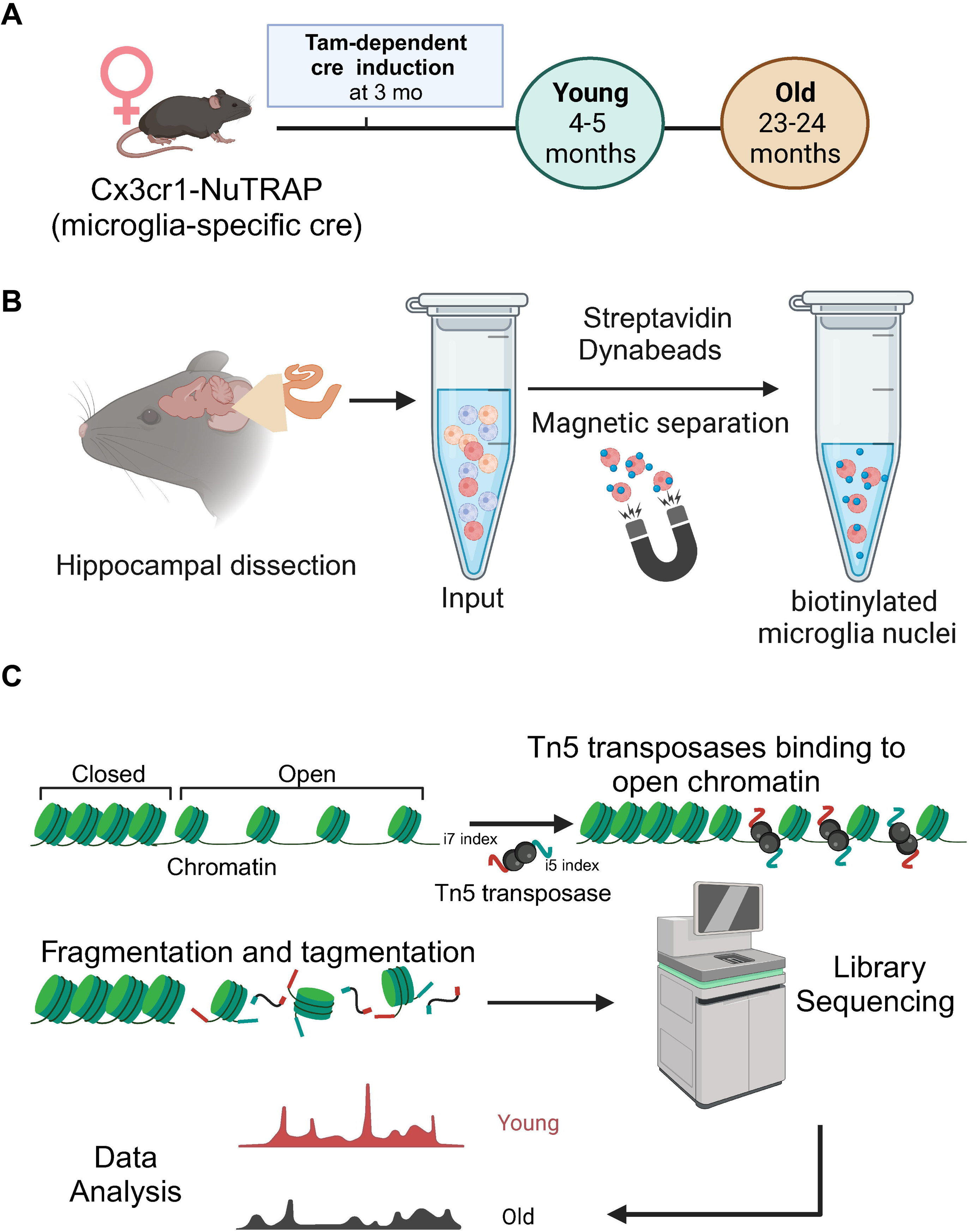
Experimental design in the study. A) Schematic of experimental group of Cx3cr1-NuTRAP mice. Mice were collected at young (4-5 months) and old (23-24 months) timepoints for two groups after cre-mediated labeling at 3 months. B) Hippocampal microglia nuclei were isolated using the Isolation of Nuclei TAgged in specific Cell Types (INTACT). C) The Assay for Transposable Accessible Chromatin using Sequencing (ATAC-Seq) libraries were prepared using approximately 50,000 nuclei per sample. Hyperactive Tn5 transposase was used to probe DNA accessible regions, inserting sequencing adapters into accessible chromatin areas. Purified libraries were sized, quantified, and subsequently subjected to sequencing. The resulting reads were aligned to the mm10 genome, filtered, deduplicated and subsampled. Broad (401bp) ATAC peaks were called using MACS2, regions with increased or decreased accessibility were identified and mapped to genomic features, locations and transcription factor binding sites.

## 2. Materials and Methods

### 2.1. Experimental Animals

All procedures involving mice were approved by the Institutional Animal Care and Use Committee (IACUC) of Oklahoma Medical Research Foundation (OMRF) and were carried out in accordance with the National institutes of Health (NIH) Guide for the Care and Use of Laboratory Animals. Mice were bred and housed in the animal facility at OMRF, maintaining specific pathogen-free conditions within a HEPA barrier environment. Breeder mice were obtained from Jackson Laboratories (Bar Harbor, ME). Cx3cr1-Cre/ERT2^+/+^ males (stock #020940) were mated with NuTRAP^flox/flox^ females (stock #029899) to generate the desired Cx3cr1-cre/ERT2^+/wt^; NuTRAP^flox/wt^ (Cx3cr1-NuTRAP mice) progeny, as previously described ^20^. In Cx3cr1-NuTRAP mice, upon cre recombination in Cx3cr1-expressing cells, there is the deletion of the floxed stop cassette, resulting in the activation of the NuTRAP allele ^20,23^. This activation enables the labeling of microglial ribosomes with eGFP, while the nuclei are labeled with biotin and mCherry ^23^. Female Cx3cr1-NuTRAP mice were allowed to age to 4-5 months and 23-24 months before tissue collection (n = 4 samples/age) **(Figure 1A)**. DNA was extracted from mouse ear punch samples for genotyping. Genotyping was performed using standard PCR detection of Cx3cr1-cre/ERT2 (Jax protocol 27232; primers: 20669, 21058, 21059) and NuTRAP floxed allele (Jax protocol 21509; primers: 21306, 24493, 32625, 32626), as previously described ^20^. Mice were euthanized by cervical dislocation, followed by rapid decapitation, in accordance with the AVMA Guidelines for the Euthanasia of Animals.

### 2.2. Tamoxifen (Tam) induction of cre recombinase

At approximately 3 months of age, mice were given intraperitoneal injections of tamoxifen (Tam) solubilized in 100% sunflower seed oil by sonication (100 mg/kg body weight, 20 mg/ml stock solution, #T5648; MilliporeSigma) daily for five consecutive days, as previously described ^20,23,24^. For the chromatin accessibility experiments described in this study, hippocampal tissues were collected from mice aged 4-5 months and 23-24 months. This timeframe allowed for a minimum of one month following the last Tam injection, ensuring the turnover of Cx3cr1-expressing circulating cells ^25^.

### 2.3. Isolation of Cx3cr1-NuTRAP hippocampal microglia nuclei

Hippocampi were collected from female Cx3cr1-NuTRAP mice at ages 4-5 months and 23-24 months and rinsed in 1× Phosphate Buffered Saline (PBS). For each ATAC-Seq sample, hippocampi from 2 age-matched mice were pooled (4 hippocampi per sample for an n = 4 samples per age). For each sample, hippocampi were minced into small pieces and homogenized in 4 ml ice-cold nuclei EZ lysis buffer (#NUC-101, Millipore Sigma) supplemented with 1×Halt protease inhibitor cocktail (ThermoFisher Scientific) using a glass Dounce tissue grinder set (#D9063; Millipore Sigma: 20 times with pestle A and 10 times with pestle B). To release any remaining sample from pestles, Pestles A and B were washed with 50 µl of ice-cold nuclei EZ lysis buffer containing the protease inhibitor cocktail. Undissociated tissues were removed by centrifugation at 200 × g for 1.5 minutes at 4°C. The resulting supernatant, containing the nuclear material, was filtered through a 30 µm wet cell strainer and subsequently centrifuged at 500 × g for 5 minutes at 4°C. The pellet containing the nuclei was vortexed briefly and resuspended by vortexing again in 250 µl nuclei EZ lysis buffer at a moderate to high speed. Subsequently, 1.7 ml of nuclei lysis EZ buffer was added, and the suspension was incubated on ice for 5 minutes. The nuclei suspension was then centrifuged at 500 x g for 5 minutes at 4°C.

The resulting nuclei pellet was resuspended in 200 µl ice-cold nuclei EZ storage buffer and diluted with 1.7 ml nuclei purification buffer (NPB: 20 mM HEPES, 40 mM NaCl, 90 mM KCl, 2 mM EDTA, 0.5 mM EGTA, 1x Halt protease inhibitor cocktail), and subjected to the Isolation of Nuclei TAgged in specific Cell Types (INTACT) protocol ^20,23^. Briefly, 50 µl of resuspended M-280 Streptavidin Dynabeads (#11205, ThermoFisher Scientific) were added into a fresh 2 ml microcentrifuge tube. The Dynabeads were washed three times with 1 ml of NPB using a DynaMag-2 magnet (#12321; ThermoFisher Scientific), with each wash performed after 1 minute of incubation. After washing, the beads were reconstituted with NPB to their initial volume of 50 µl and then gently mixed with the nuclear suspension. The mixture of nuclei and magnetic beads was incubated at 4°C for 40 minutes, using gentle rotation settings, to facilitate the affinity binding of streptavidin beads to the cell-specific, biotinylated nuclei. After incubation, the streptavidin-bound nuclei were magnetically separated with the DynaMag-2 magnet and beads were resuspended in 50 µl of ice-cold NPB **(Figure 1B)**.

### 2.4. Library construction and Assay for Transposase-Accessible Chromatin using sequencing (ATAC-seq)

A 3 µl-volume of the streptavidin-bound nuclei was diluted with 10 µl of nuclease-free water, followed by staining with 10 µl of Trypan blue and counted using a Countess 2 Automated Cell Counter (ThermoFisher Scientific). The total nuclei count was adjusted by multiplying it with the dilution factor of 3.33. ATAC-seq libraries were prepared using a volume of the streptavidin-bound nuclei suspension equivalent to approximately 50,000 nuclei, according to the protocol provided by the manufacturer and previously described ^21,22^. Using the Illumina Tagment DNA TDE1 Enzyme and Buffer Kit, the transposition reaction mixture (25 µl TD – 2x reaction buffer, 2.5 µl TDE1-Nextera Tn5 transposase and 22.5 µl of nuclease-free water) was mixed with the streptavidin-bound nuclei suspension and incubated at 37°C for 30 minutes.

Following the transposition reaction, transposed DNA were purified using the Qiagen MinElute PCR Purification Kit. Purified transposed DNA were initially PCR amplified for 5 cycles using Nextera Indexes (Nextera XT Index Kit v2–Set A). To reduce GC size bias, qPCR was performed to determine the appropriate number of cycles to amplify DNA prior to saturation. The transposed DNA were further PCR amplified for the number of cycles corresponding to ¼ of the maximum fluorescent intensity detected during qPCR (∼5-6 cycles). The amplified libraries were purified using Qiagen MinElute PCR Purification Kit and eluted in 20 μL. Libraries were sized and quantified using TapeStation High Sensitivity DNA ScreenTape (Agilent Technologies). Libraries for each sample were normalized and pooled at a concentration of 4 nM, denatured, and diluted to 12 pM for sequencing on the NovaSeq 6000 system (SP, PE50bp and PE75bp, Illumina) according to the manufacturers guidelines **(Figure 1C)**.

### 2.5. ATAC-seq data analysis

Following sequencing, paired-end reads were checked for quality using fastQC ^26^, and adaptor-trimmed using Trimmomatic ^27^ v0.35. End-trimming removed leading and trailing bases with a Q-score<25, cropped 5 bases from the start of the read, dropped reads less than 50 bases long, and dropped reads with average Q-score<25. Alignment was performed using Bowtie 2 ^28^ (v2.3.1) against the mouse reference genome (GRCm38/mm10), using-X 2000 to allow for reads separated by <2kb. Files were converted to Binary alignment map (BAM) files and reads arising from PCR duplicates were removed using Samtools ^29^. Reads were then sorted, indexed, reads mapping to the mitochondrial genome and chromosome Y removed, and converted to Browser extensible data (BED) files using Samtools ^29^ (v1.14). Broad peak calling was carried out using MACS2 ^30^ (v2.2.7.1) with a 200bp smoothing window and an offset of 100. As we were calling broad peaks, reads were not shifted + 4 bp and − 5 bp for positive and negative strand respectively, to account for the 9-bp duplication created by DNA repair of the nick by Tn5 transposase and achieve base-pair resolution of TF footprint analyses. Additionally, fragment size distribution and nucleosome positioning of ATAC-seq libraries were assessed using ATACseqQC^31^. Reads were randomly subsampled for an approximate 38 million reads per sample **(Additional file S1)**.

The R package, DiffBind ^32^ was used to quantify the read counts, and peaks were normalized to the number of reads. A set of consensus peaks was created with peaks that were identified in at least three samples from each group. Differentially accessible regions (DARs) (p-value ≤0.05 and False discovery rate ≤0.01) were identified using DESeq2 ^33^ from comparisons between the young (4-5 months) and old (23-24 months) samples. DiffBind ^32^ was used to generate the plots for Principal Component Analysis (PCA), and Volcanos of DARs. Circos plots and feature/density distribution analyses were made in R using Circlize ^34^ and ChIPseeker ^35^, respectively. Statistical analysis, including log odds ratios was performed using Microsoft Excel and GraphPad Prism 9 (San Diego, CA). Gene ontology analysis including the dot plot and CNET plot was performed in R using the enrichGO ^36^ package from clusterProfiler ^37^. DARs peak files were analyzed for known and *de novo* motifs enrichments using HOMER ^38^. Identifying gene enrichments closer to DARs were performed by using bedtools ^39^ to annotate sets of loci with features from UCSC. Hypergeometric test was used to perform a comparative analysis to assess the overlap of DARs identified within our study and those from other studies.

## 3. Results

Following ATAC-seq, an average of 38 million reads successfully passed quality control per sample (Additional file 6). After alignment, filtering, removal of reads mapped to the mitochondrial genome, and removal of duplicate reads, remaining reads were randomly subsampled to equal numbers for each sample. To ensure consistency between the young and old samples for quality control post-alignment, the distribution of fragment sizes was assessed. The fragment distribution plots show clear peaks corresponding to nucleosome-free regions (NFR) with sizes below 100 bp, as well as peaks for mono- (∼200 bp), di- (∼400 bp), and tri- (∼600 bp) nucleosomes, similar in both the select young and old samples **(Figure S1A)**. As expected, a high proportion of the ATAC-seq libraries contain short reads (∼100bp) from nucleosome-free regions **(Figure S1A)**, indicating high quality of the libraries since the Tn5 transposase prefers more accessible chromatin regions ^22,31,40–42^. The presence of mono-nucleosome fragments **(Figure S1A)** represent nucleosome-bound open chromatin regions, as the Tn5 transposase also inserts sequencing adapters into the linker DNA between adjacent nucleosomes ^31^ **(Figure 1C)**. Further quality control measures were carried out to assess the position of nucleosomes around the TSS by analyzing reads aligned to chromosome 1 in the select young and old samples. A plot of the nucleosome positioning shows an enrichment of NFRs precisely at the transcription start site (TSS) while nucleosome-bound regions are depleted near the TSS with an increasing enrichment away from the TSS **(Figure S1B)**, typical of ATAC-seq libraries ^31,40,42^.

### 3.1 Overview of microglia chromatin accessibility landscape

In total, 138,421 open chromatin regions or peaks were identified across all hippocampal microglia samples **(Additional file S2)**. Approximately 14.73 % of these peaks were located within 1kb of the transcription start site while a majority of peaks were found within intronic and exonic intergenic regions or in intergenic regions **(Figure S2A)**. The proportion of peaks in promoter-flanking regions and exons was lower, indicating less accessible chromatin in comparison to the regions around the TSS, intergenic, and intronic regions **(Figure S2A & S3)**. Generally, functional regions such as promoters tend to have lower nucleosome occupancy, whereas most genomic DNA exhibits a higher nucleosome density ^21,40,42,43^.

Subsequently, a consensus set of 53,421 peaks was created using an inclusion criterion that required a peak to be in at least three samples within each age group **(Figure 2A)**. Of the identified consensus peaks, 26,191 (49.03%) were shared between young and old samples while 22,276 (41.70%) and 4,954 (9.27%) peaks were unique to young and old samples, respectively **(Figure 2A & B**; **Additional file S3)**. Using the consensus set of peaks, a separation of the young and old samples in the first component of a Principal Component Analysis plot was evident **(Figure 2C)**, demonstrating distinct patterns of chromatin accessibility with age. Overall, there were more peaks identified in young hippocampal microglia samples compared to the old **(Figure 2A & B)**. In contrast, a recent finding by Li et al. ^11^, suggests an increase in the number of ATAC peaks in aged female microglia. It is worth noting that, in their study, microglia were isolated from whole brain, excluding the cerebellum ^11^. Microglia populations are known to exhibit regional variations within the brain, particularly in response to perturbations such as aging ^44,45^. This distinction could potentially account for the differences in the ATAC peaks observed, along with differences in the method of peak calling.

**Figure 2.**
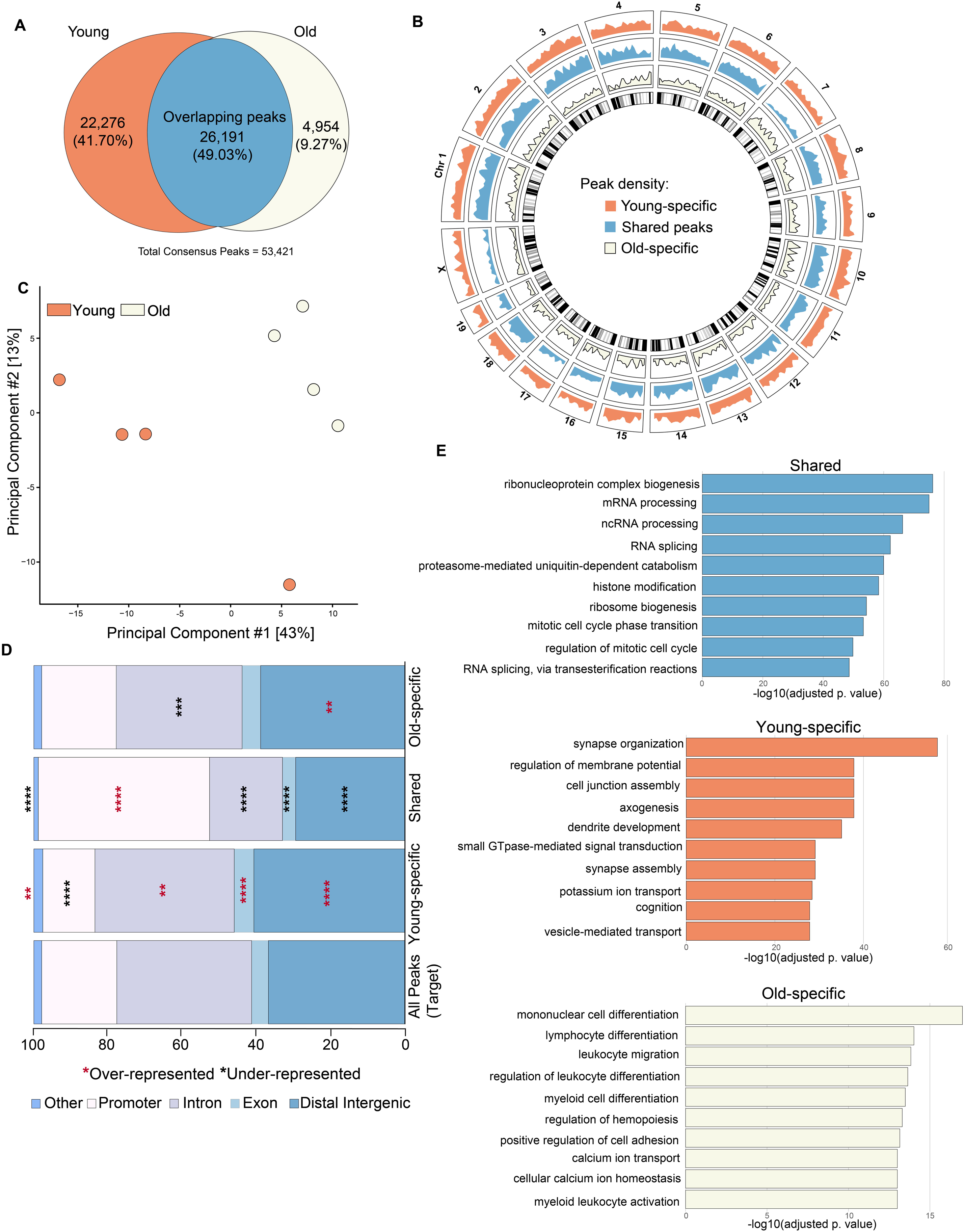
Overview and landscape of chromatin accessibility with age. A) A set of consensus peaks totaling 53,421 was identified, with each peak appearing in at least 3 samples in at least one group. Within this set, 26,191 peaks (representing 49.03%) were shared between the young and old groups, 22,276 peaks (41.70%) were exclusive to the young, and 4,954 peaks (9.27%) were exclusive to the old. B) Peaks were distributed across the genome where the density was notably higher in the young-specific and shared peaks sets than in the old-specific group. C) A principal component analysis performed on the entire set of consensus peaks (53,421) identified revealed a distinct separation between young and old samples in the first component. Notably, the young samples exhibited higher variability, compared to the old. D) The annotation of enrichment patterns for ATAC peaks, whether shared, young-specific, or old-specific, was performed in relation to the entire set of 138,421 peaks (target). The plot represents the distribution of ATAC peaks in promoters, exons, introns, distal intergenic, and other regions (3’ UTR, 5’ UTR, and Downstream (≤300bp)). Young-specific peaks were over-represented in all annotated regions except in promoters, where they were underrepresented. Shared peaks were over-represented exclusively in promoters and under-represented in all other regions. Old-specific peaks were under-represented in distal intergenic regions but under-represented in introns (*p < 0.05, **p < 0.01, ***p < 0.001, ****p < 0.0001, Fisher’s exact test, blue stars indicate over-representation, while black stars denote under-representation). E) ATAC peaks were annotated to the nearest genes within a range of ±3kb, and GO enrichment analyses were performed on the gene sets corresponding to shared, young-specific, and old-specific peaks. The plots show the Top 10 GO Biological Processes for genes closest to the various sets of peaks (adjusted p value < 0.05).

### 3.2 Feature distribution and biological processes associated with distinct and shared peaks

Next, we used ChIPseeker ^35^ and ChIPpeakAnno ^46^ to examine the distribution of consensus peaks, whether shared or specific to young or old microglia, across different genomic features, with respect to a background of all the identified peaks (138,421 peaks). Shared peaks between old and young hippocampal microglia were predominantly enriched in promoter regions **(Figure 2D)**. In contrast, peaks specific to young samples were overrepresented in introns, exons and other regions while being depleted in promoter regions **(Figure 2D)**. Peaks specific to old samples were overrepresented in intergenic regions but underrepresented in intronic regions **(Figure 2D)**. This distinct distribution of peaks suggests the presence of both shared and age-associated mechanisms of chromatin regulation in female microglia, possibly associated with diverse biological functions and processes.

To provide biological contexts to shared and age-specific peaks, the peak sets were annotated to the nearest genes (within ±3kb) and subsequent biological pathway analyses were performed using Gene Ontology (GO) ^36,37^. Analysis of biological function of genes nearest to shared peaks revealed that these genes are mainly associated with pathways involved in the post-transcriptional regulation of gene expression, RNA metabolism and chromatin remodeling. These pathways are integral to gene regulation and normal cellular processes which are fundamental to both old and young microglia ^47–50^ **(Figure 2E)**. Genes nearest to peaks specific to young microglia were associated with pathways critical for establishing neural circuits and neuronal communications such as synaptic organization, regulation of membrane potential and neuronal axon generation **(Figure 2E)**. On the contrary, genes nearest to peaks specific to old were linked to pathways such as myeloid cell differentiation essential to immune system maintenance and response to stimuli **(Figure 2E)**.

### 3.3 Gains and losses of chromatin accessibility with aging

To investigate the changes in chromatin accessibility with age in female mice, we compared the accessibility of young and old microglia. Before calling differentially accessible regions (DARs) between young and old, the set of consensus peaks were re-centered (width = 401bp) and peaks within 400bp of each other were merged resulting in a total of 48,353 peaks **(Additional file S4)**. We identified 1270 (2.6%) regions that were differentially accessible between young and old female microglia (DESeq2, FDR ≤0.010) **(Figure S2B; Additional file S4)**. DARs show changes in chromatin accessibility (Old vs Young) in older animals with both increases and decreases observed **(Figure 3A)**. The majority of DARs, 1077 peaks representing 84.80% of identified regions, increased accessibility in the old microglia whereas 193 (15.20%) regions decreased in accessibility in old microglia **(Figure S2C & S2D)**. These findings suggest a degree of conservation of chromatin accessibility with age, as only a limited number of regions significantly changed in accessibility within the hippocampal microglia of female mice. Furthermore, there is a tendency for these regions to become more accessible rather than losing accessibility.

**Figure 3.**
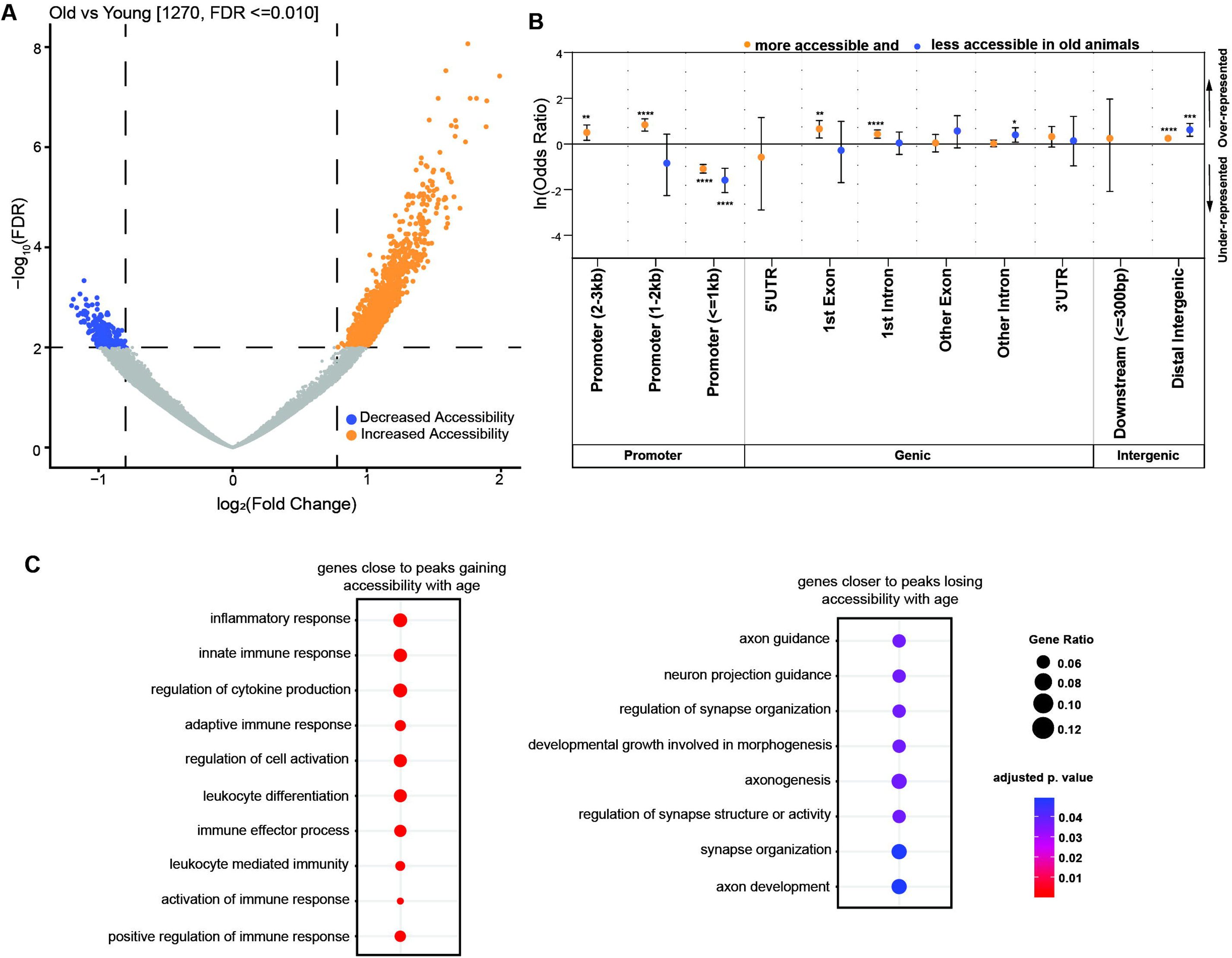
Differential chromatin accessibility changes with age. The set of consensus peaks were re-centered (width = 401bp), and peaks within 400bp proximity were merged, yielding a total of 48,353 peaks before the identification of differentially accessible regions (DARs) between young and old samples. We detected 1,270 regions (2.63%) that exhibited differential accessibility in female microglia between the young and old groups (DESeq2, FDR ≤ 0.010, p-value ≤ 0.05). A) The volcano plot of DARs depicts a higher number of regions increasing in accessibility with aging (orange) compared to those decreasing in accessibility (blue). B) The natural logarithm of the Odds ratio was computed for DARs across genomic features through ChIPSeeker analysis. Enrichment comparisons were conducted for regions with increased (orange) and decreased (blue) accessibility. Odds ratios exceeding 1.0 were deemed over-represented, while those below 1.0 were considered under-represented. Significant enrichment or depletion is indicated by stars (*p < 0.05, **p < 0.01, ***p < 0.001, ****p < 0.0001, Fisher’s exact test). C) DARs were annotated with the nearest genes (±3kb), and GO enrichment analyses were carried out on the gene sets linked to peaks exhibiting increased accessibility (595 genes) or decreased accessibility (93 genes). The dot plot illustrates the top 10 GO Biological processes.

To increase the rigor of our study, we performed a comparative analysis of DARs identified within our study and in the study by Li et al ^11^. This study examined the impact of lipopolysaccharide (LPS) treatment on chromatin accessibility with age in microglia isolated from the whole brain (excluding the cerebellum) of female mice by flow cytometry. As a control, a separate cohort of female mice were treated with Phosphate-buffered saline (PBS). Using the DAR identification pipeline described in this study, we identified DARs in microglia isolated from young (3 months) and old (24 months) PBS-treated mice (n = 2 per age group), mirroring the age groups in our study. A total of 80,267 consensus peaks were identified **(Figure S4A)**, with the inclusion criterion that a peak was present in both samples within each age group, since there were only two samples per age group. Among the identified consensus peaks, 1477 were differentially accessible **(Additional file S5)**. Of these consensus peaks, 1475 showed increased accessibility, while two (2) regions, decreased accessibility in the older group. To compare our findings with those of Li et al., we performed an overlap and intersection analyses of DARs across both studies. Our analyses revealed a total of 79 DARs across the studies found within 10kb of each other **(Figure S4B)**, with 38 overlapping DARs (**Figure S4C)**. Despite differences in experimental design, including brain region used for microglia isolation, the observed commonalities between the studies were greater than expected by random chance. The common peaks were nearly universally areas of increased accessibility in both studies.

### 3.4 Differential accessibility is underrepresented at the proximal promoter in female microglia with age

To understand how the changes in chromatin accessibility may affect the gene regulation, we examined DARs to identify whether they are overrepresented or underrepresented within specific genomic locations when comparing young and old female microglia. Proximal promoters (≤1kb), while containing approximately one-third (27.66% - 13,375) of the consensus peaks **(Figure S2B)**, were underrepresented for DARs **(Figure 3B)**. In contrast, distal promoter regions (1-3kb) were enriched with regions that gained accessibility compared to regions that decreased accessibility **(Figure 3B)**. Further, DARs demonstrated an enrichment in distal intergenic regions regardless of whether they were gaining or losing accessibility, whereas gains in accessibility were overrepresented in the first introns and exons **(Figure 3B)**. Studies have reported that the chromatin landscape of enhancers and promoters define the identities of tissue-resident macrophages ^51,52^. Our findings indicate that while female microglia can indeed respond to the aging process by altering chromatin accessibility, these modifications are depleted in the proximal promoter, a key regulatory region for microglia gene expression and genomic accessibility. Instead, it appears that changes in chromatin accessibility may be primarily regulated distally to gene bodies, through other functional elements such as enhancers in non-promoter regions. Still, even minimal changes in promoter accessibility may have a crucial role in impacting microglia function. Interrogation of these data with hippocampal microglia-specific RNA sequencing data in young and old females may aid in identifying whether small changes in the chromatin architecture impact gene expression changes with age.

### 3.5 Biological processes associated with age-associated chromatin accessibility changes

To further contextualize the set of regions that gained or decreased accessibility with aging, we identified the closest genes (within ±3kb of genes) to DARs and performed functional analysis using clusterProfiler ^37^. Nearest genes (±3kb) to the set of all consensus peaks were used as background (target) **(Additional file S6)**. Genes near regions with gains in accessibility were linked to inflammatory pathways and processes **(Figure 3C)**. Among the top Gene Ontology (GO) ^36^ biological processes were inflammatory response, innate and adaptive immune responses, as well as the regulation of cytokines and cell activation **(Figure 3C)**. This is consistent with the annotated closest genes such as Toll-like Receptors (TLRs), such as *TLR-4* **(Figure S5)**, involved in innate immune recognition ^53,54^ and associated with neuroinflammation and neurodegeneration ^55,56^. Other annotated genes include Interferon Regulatory Factor 5 (*IRF5*) and AXL receptor tyrosine kinase (*AXL*), a member of the Tyro3-Axl-Mer receptor family of tyrosine kinases ^57^ **(Figure S5)**. *IRF5* is crucial in the expression of proinflammatory cytokines such as Interleukin-6 and the regulation downstream immune response by TLRs ^58,59^. *AXL* expression is increased during inflammation and disease ^57,60^, and it is linked to phagocytosis ^60–62^, regulation of inflammatory response and microglial survival ^61^.

Conversely, genes annotated to regions that decreased accessibility were linked to genes including like Gamma-aminobutyric acid (*GABA*) receptor subunit alpha-2, which is reported to be expressed in microglia cell lines ^63,64^, and Myosin VB (*MYO5B*) **(Figure S6)**, whose expression decreases in microglia following intraperitoneal injection of LPS in mice ^65^. GO analysis of these genes revealed pathways and processes associated with axon guidance and synaptic function **(Figure 3C)**. Taken together, these data show that increases in chromatin accessibility in female microglia are mainly located near genes involved in microglia activation and inflammation. In contrast, decreasing chromatin accessibility is primarily associated with microglial genes related to neuronal and synaptic function. This is in line with gene expression changes that have been observed in female hippocampal microglia with aging, which points to alterations in the chromatin landscape as a potential driver for these gene expression changes ^12^.

### 3.6 Age-associated changes in gene regulatory networks

Given that open chromatin affects transcription by facilitating transcription factor (TF) binding, we analyzed the enrichment of TF binding motifs in regions of increased or decreased accessibility using Homer ^38^. Motif enrichment analyses revealed *de novo* and known consensus sequences enriched in regions that increased or decreased accessibility in old microglia **(Additional file S7-S10)**. In regions with increased accessibility, we observed a significant enrichment of transcription factor binding sites including members of the E26 transforming sequence (ETS) family of TFs such as PU.1 (encoded by *SPI1*) ^66^ and the E74 Like ETS transcription Factor 5 (ELF5) **(Figure 4A)**. The binding of PU.1 is known to initiate nucleosome remodeling and H3K4 mono-methylation, potentially leading to increased accessibility of chromatin or the presence of enhancers impacting the expression of genes associated with microglia activation and function ^67,68^. Additionally, PU.1 binding is reported to affect inflammatory response of microglia to stimuli like Lipopolysaccharide ^69^. The *de novo* enrichment analysis revealed best matched TFs including Interferon Regulatory Factor 8 (IRF8 and SMAD family member 2 (SMAD2). These transcription factors play critical roles in myeloid cell differentiation as well as microglial activation ^70–73^. These findings suggest that alterations in chromatin accessibility at TF binding sites for DNA-binding proteins such as PU.1 may control microglia inflammatory response to age in female mice.

**Figure 4.**
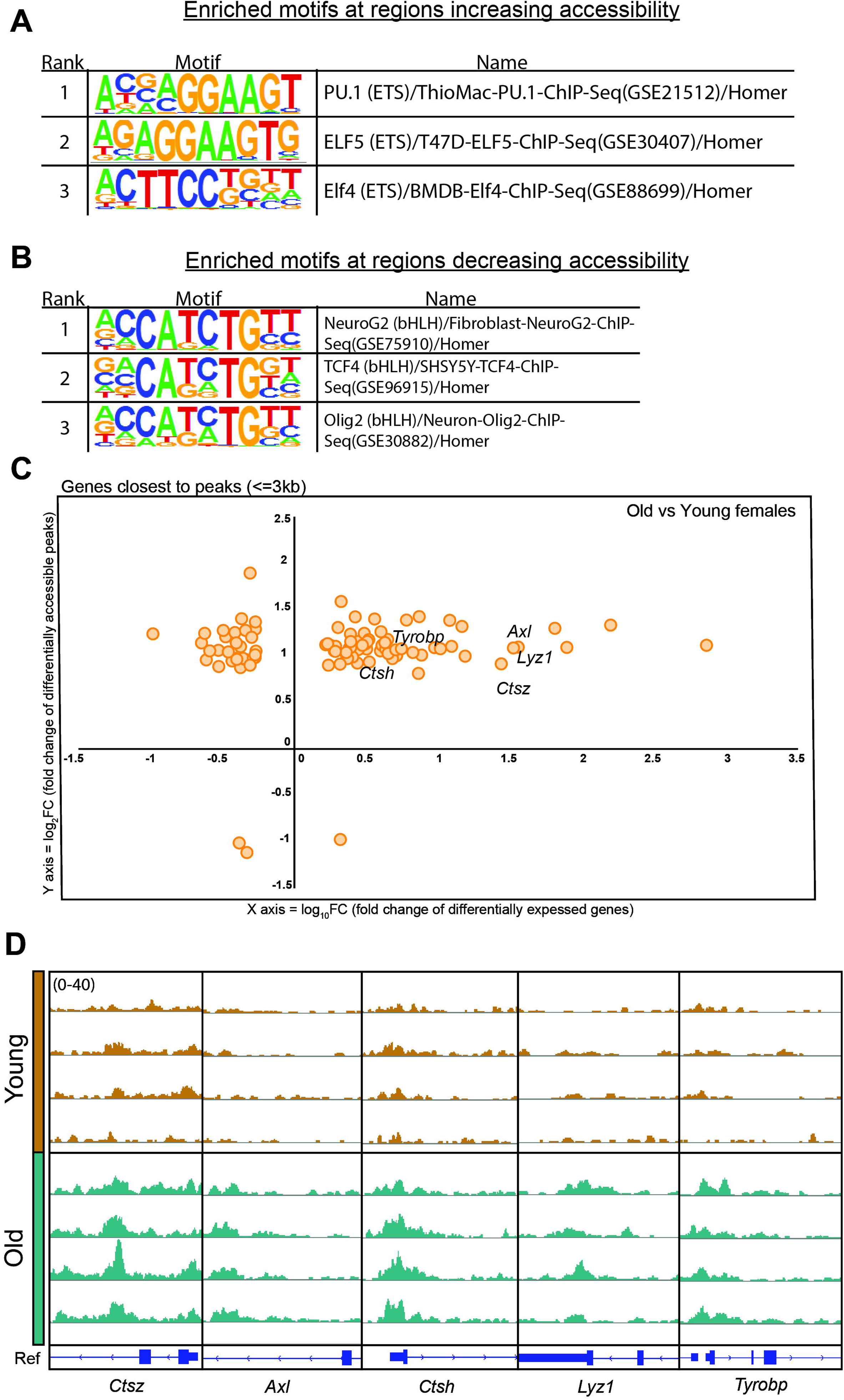
Motif enrichment and correlation with gene expression. Tables displaying enriched motifs and their corresponding DNA-binding proteins as identified by HOMER analysis for regions that A) increased in accessibility (highlighting motifs associated with myeloid cell differentiation and function) and B) decreased accessibility (featuring motifs linked to neuronal cell differentiation and the activation of myelin-associated gene expression) in the old group. C) Comparison of the nearest genes annotated to differentially accessible regions (DARs) with the differentially expressed genes (DEGs) identified in a prior study conducted by our group on young and old female microglia. In the study by Ocañas et al. ^12^, the effects of sex on the translatome were investigated in hippocampal microglia of both young (5 – 6 months) and old (22 – 25 months) mice, using Cx3cr1-cre/ERT2 (Jung) mice. The plot depicts the correlation between differential accessibility and gene expression (log (FC)) in female microglia.

Conversely, in regions with decreased accessibility, we found enriched TF motifs for Oligodendrocyte Transcription Factor 2 (OLIG2), neurogenic differentiation 1 (NEUROD1), and neurogenin 2 (NEUROG2), associated with neurogenesis and neuronal differentiation ^74–78^ **(Figure 4B)**. Studies report that transformation of microglia and astrocytes is possible through the expression of NEUROD1, which is associated with neuronal fate and identity ^79,80^. Overall, we observed differential accessibility at TF binding motifs that have downstream functions in inflammation and neuronal differentiation, aligning with pathways enriched in DARs.

### 3.7 Relationship of chromatin accessibility to gene expression

We next compared the genes closest to DARs to differentially expressed genes (DEGs) between young and old female microglia in a previous study from our group. Ocañas et al. ^12^, studied sex effects on the translatome and transcriptome of both young (5 – 6 months) and old (22 – 25 months) hippocampal microglia. We sought to draw comparisons and correlate chromatin accessibility to gene expression observed between young and old female microglia. It is worth noting that Ocañas et al. ^12^, used the Cx3cr1-cre/ERT2 (Jung) – NuTRAP line in their study, while this study uses the Cx3cr1-cre/ERT2 (Litt) – NuTRAP line. We compared the 687 genes annotated to DARs (whether increasing or decreasing accessibility) to the 1190 DEGs identified **(Additional file S11)** ^12^,and found 94 genes in common. Subsequently, the log_2_ accessibility fold change was plotted against the log_10_ expression fold change for these 94 genes **(Figure 4C)**. Interestingly, we did not observe a clear-cut positive correlation between the extent of chromatin accessibility and gene expression **(Figure 4C)**. Our data points were distributed across four quadrants, with majority of data points in the quadrant representing an increase in chromatin accessibility alongside an increase in gene expression **(Figure 4C)**. The data points in this quadrant included genes that increase in expression with age like *AXL*, cathepsin Z, and H (*CTSZ* and *CTSH*), which exhibited an increase in chromatin accessibility **(Figure 4D)**. While it is possible that the differences in mouse lines used in the two studies contribute to the limited correlation, these findings underlie the complex nature of gene expression regulation, which may involve additional epigenetic mechanisms such as DNA methylation as well as more complex and distal correlation through enhancers.

### 3.8 Differential chromatin accessibility on the X chromosome of aging female microglia

We examined the chromosomal location of DARs and observed changes in chromatin accessibility across all chromosomes, showing varying proportions of regions gaining or losing accessibility. Interestingly, chromosomes 7 and 17 lacked regions with decreased accessibility, and chromosomes 12, 16, and X had fewer regions showing decreased accessibility **(Figure 5A)**. Early in mammalian development, inactivation of one X chromosome (XCI) occurs in females to dosage compensate X encoded genes ^81^. Following development, this inactivation (Xi) is maintained through repressive epigenetic mechanisms such as hypermethylation and heterochromatin ^82^. Escape from X inactivation has been proposed to occur with aging in females ^83,84^. Focusing on Chromosome X, we examined alterations in its chromatin accessibility independently of autosomal accessibility. We segregated X chromosome peaks from autosomal peaks, re-centered peaks, and re-analyzed to identify DARs found within X chromosome. A consensus set of 3,579 Chromosome X peaks was identified using an inclusion criterion that required a peak to be in at least three samples within each age group, as described previously.

**Figure 5.**
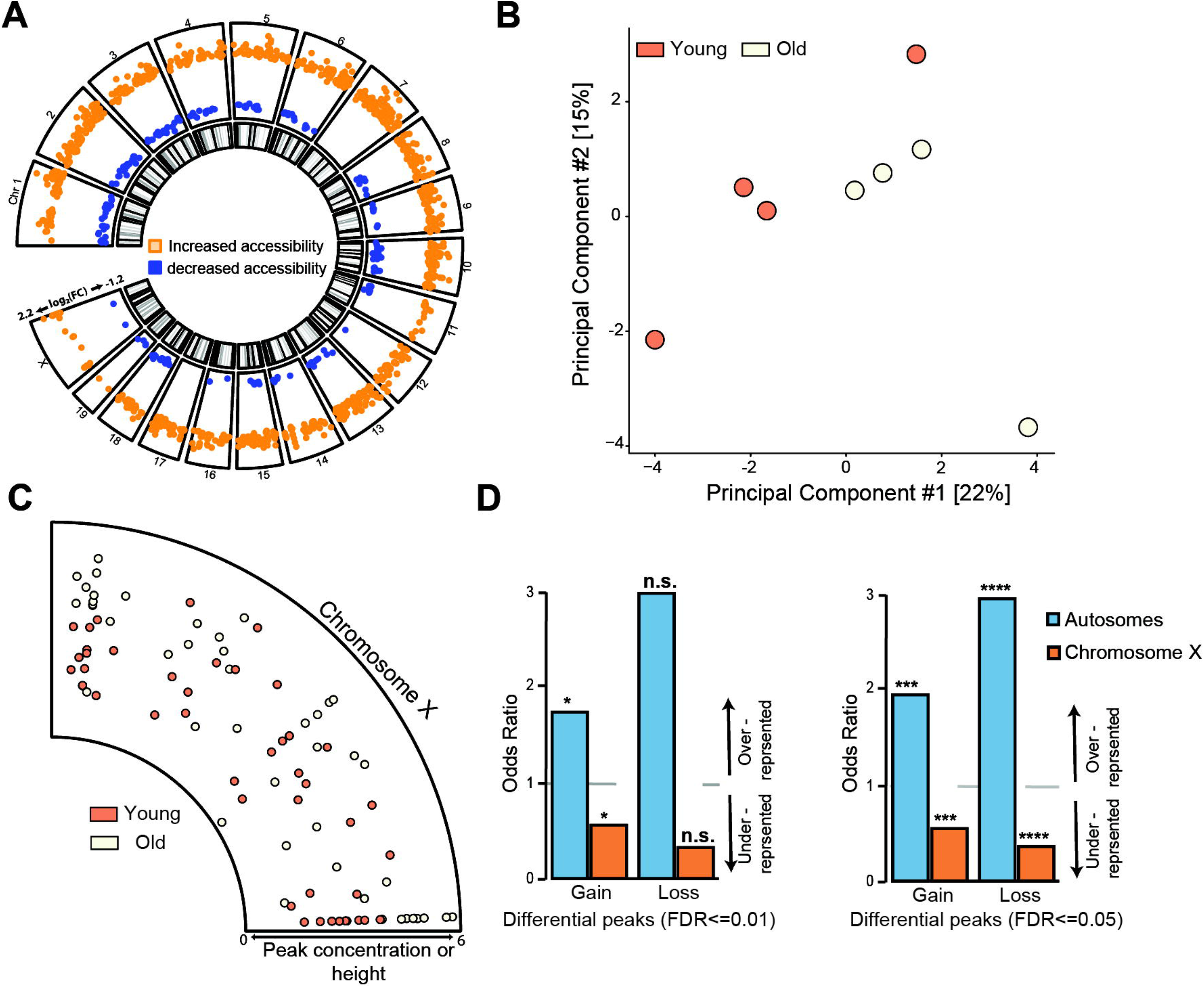
X chromosome chromatin accessibility. A) Circos plot depicting DARs across chromosomes reveals a consistent trend where regions that change with aging tend to increase in accessibility than lose accessibility. B) X chromosome peaks were separated from autosomal peaks, and a re-analysis was conducted to identify DARs within chromosome X. After re-centering the peaks, a consensus set of 3,579 peaks present in at least three samples within at least one age group were identified on Chromosome X. Principal Component Analysis X chromosome consensus set of peaks demonstrates a clear separation between the young and old samples in the first component based on age. C) A Circos plot visualizing the concentration (peak height) in both young and old samples across the 45 regions identified as changing in accessibility with aging on the X chromosome (FDR ≤ 0.05, p-value ≤ 0.05). D) Odds ratios were calculated to assess the enrichment of DARs on Chromosome X compared to autosomes through ChIPSeeker analysis. Enrichment comparisons were made for regions that either increased (Gain) accessibility and those that decreased (Loss) accessibility, using an FDR of ≤ 0.01 or ≤ 0.05. Odds ratios exceeding 1.0 denote over-representation, while ratios below 1.0 indicate under-representation. The significance of enrichment or depletion is indicated by stars (*p < 0.05, **p < 0.01, ***p < 0.001, ****p < 0.0001, Fisher’s exact test). We observe fewer changes on Chromosome X than would expected by chance.

Using the consensus peaks, a separation of the young and old samples in the first component in a Principal Component Analysis was evident **(Figure 5A)**. Due to the limited number of peaks on Chromosome X, we identified DARs with an FDR cutoff of 0.050 (DESeq2, p-value ≤0.05), resulting in the detection of 45 X chromosome DARs **(Figure 5B; Additional file S12)**. As identified with the global chromatin accessibility changes, a majority of DARs (42 DARs - 73.3%) were found to increase in accessibility **(Figure 5C)**. Finally, we examined the likelihood of DARs occurring on Chromosome X in comparison to the autosomes. There was an underrepresentation of DARs on chromosome X **(Figure 5D)** compared to autosome. This underrepresentation of differential accessibility on the X chromosome was observed regardless of whether regions were gaining or losing accessibility in aged microglia. The data suggest that age-related accessibility changes occur in a chromosome-independent manner, resulting in increased microglia chromatin accessibility with age. This study is limited regarding the separation of the active (Xa) and inactive X (Xi), thus we are unable to distinguish the specific contributions of Xa or Xi to chromatin accessibility changes with age. Allele-specific ATAC-seq would be required to thoroughly study age-associated changes in chromatin accessibility on the X chromosome, which is beyond the scope of the current study. Nevertheless, these data do not strongly support increased escape from X inactivation with aging.

## 4 Discussion

Here, we profile the changes in chromatin accessibility of hippocampal microglia in female mice with aging. Understanding the genome regulatory landscape of microglia in aging is crucial for examining reactivational microglial states observed with aging, as well as developing strategies to delay or reverse age-related microglial dysfunction. Our study revealed: (1) female microglia respond to aging through alterations in chromatin accessibility; (2) these changes tend to result in increased accessibility rather than decreased accessibility; (3) proximal promoter chromatin accessibility is relatively stable with age, suggesting that accessibility of other distal regulatory elements plays a greater role in the regulation of microglial aging; (4) regions of increased accessibility are located near genes involved in immune and inflammatory responses, while decreased accessibility are around genes involved in regulating neuronal function; (5) the potential DNA-binding proteins that regulate the functional phenotypes in aged female microglia; and (6) age-related changes in chromatin accessibility are evident across all chromosomes, with significantly fewer changes observed on the X chromosome.

The “heterochromatin loss theory” of age suggest a genome-wide trend in which chromatin accessibility increases due to the loss of heterochromatic regions with aging ^85–89^. In our initial analysis of consensus peaks, we do observe peaks that are unique to both young and old microglia. We observe distinct peaks in old microglia, likely indicating regions that were not initially accessible in young microglia. However, there was a higher number of distinct peaks in the younger group compared to the old, which implies that more euchromatic regions in the young became suppressed with age. This agrees with recent a finding suggesting that overall chromatin accessibility may not change with aging ^90^. Instead, there are changes in chromatin accessibility at specific genomic regions, with both gains and losses observed. These results strongly indicate that the regulation of age-related chromatin accessibility varies across distinct genomic regions, emphasizing the importance of conducting site-specific analyses of chromatin accessibility. The disproportionate number of unique peaks observed for both young and old may be due to the differences in genomic regions at which regulation of chromatin accessibility takes place. Unique ATAC peaks in young samples are found in a wide range of genomic regions whereas peaks distinct to old are predominantly overrepresented in distal intergenic regions, suggesting a limited regulatory domain. It is worth noting that our finding of higher ATAC peaks in young microglia is contrary to a recent study which suggests an increase in the number of ATAC peaks in aged female microglia ^11^. Our study employed the INTACT approach to isolate hippocampal microglia, whereas Li et al. ^11^ used a FACS approach to isolate microglia from the whole brain (excluding the cerebellum). Moreover, the lack of transcriptional and translational inhibitors in the cell dissociation process for FACS could introduce *ex vivo* activational confounds capable of altering the phenotypic states of microglia ^25,91^. In contrast, the INTACT microglia isolation employed here is not linked with additional *ex vivo* activational artifacts ^25^. The differences in isolation method and the brain region from which microglia were obtained could explain the variations in peak numbers. Nonetheless, there were common regions between studies indicating reproducibly altered regions of chromatin accessibility. Importantly, both their study ^11^ and ours demonstrate that female microglia can respond to aging at an epigenetic level through alterations in chromatin accessibility.

Our findings reveal a relative accessibility profile between young and old microglia, with only specific genomic loci exhibiting significant changes in accessibility. Interestingly, significant changes in chromatin accessibility at specific genomic sites during aging tend to lead to increased rather than decreased accessibility. Our functional analysis indicates a trend of microglia adopting a more inflammatory phenotype as they age, typical of aging murine brains. Our data indicate enrichment of sequences for DNA binding proteins from the ETS family of TFs including PU.1, which are critical to microglia differentiation and function ^66,70,71^. Previous transcriptomic analyses reveal a higher expression level in genes like *Spi1*, which encodes PU.1, in female microglia compared to males ^12^. Changes in accessibility affecting the transcription of these genes may significantly impact the functional phenotype of aging microglia. Increased accessibility at the binding sites of TFs known to be key regulators of microglia inflammatory phenotypes could be either a cause or reflection of a reactivational phenotype. It is known that microglia also play key roles in facilitating synaptic pruning, spine formation and synaptic maturation necessary for neuronal function ^92^. In addition, microglia promote astrocytic activation during injury and can improve the survival of mature oligodendrocytes ^93^. The observed decrease in accessibility at regions associated with microglial genes linked to these functions suggests a potential suppression of neuronal supportive functions, in addition to a shift towards a predominantly inflammatory phenotype with age.

From ATAC-seq studies, gene promoters are typically constitutively accessible across a broad range of cell types ^22,42,94–98^, a trend observed in our study where a majority of ATAC peaks were found in promoters, intronic and intergenic regions. However, we note a relatively stable pattern of chromatin accessibility, particularly in the proximal promoter, suggesting a consistent level of accessibility in regions directly upstream or downstream of genes with aging. Most regions prone to changes in chromatin accessibility are distal from the TSS. Though promoters are key regulatory elements, distal regions also contain enhancers and insulators that impact the overall chromatin landscape and gene expression patterns. The dynamic nature of chromatin accessibility across various genomic regions highlights the complexity of the regulatory mechanisms underlying age-related gene expression and the overall aging process in microglia. This intricate nature of regulation includes not just the effects of chromatin accessibility but also other epigenetic mechanism like DNA methylation and their complex interactions ^99^. Together, our genomic feature analysis aligns with prior studies, suggesting variations in the regulation of chromatin accessibility across different genomic regions.

Repressive epigenetic patterns such as hypermethylation and heterochromatin play crucial roles in the maintenance of X chromosome inactivation (XCI) and the potential escape of some genes from XCI with aging ^100,101^. Therefore, understanding the chromatin landscape within the context of the X chromosome may offer insights into the impact of sex effects on immune responses and sex biases observed in brain aging and related diseases. While our design does not distinguish between the active and inactive X chromosome, we explored alterations in the chromatin structure of the X chromosome to gain a basic understanding of its chromatin accessibility changes with age. Our data reveal that changes in chromatin accessibility are less prevalent on the X chromosome compared to autosomes which argues against a general escape from X inactivation. The bias is most likely due to the existence of the inactive X chromosome, although we are unable to discriminate the allelic contributions of the two X chromosomes to the overall X chromosome chromatin landscape within the scope of this study. In addition, we observed a pattern where regions across the genome that change in chromatin accessibility are predominantly increased, consistent with a previous report in hematopoietic stem cells ^102^. This finding suggests an extensive mechanism influencing changes in microglia phenotype with aging. Additional studies are required to mechanistically understand how chromatin accessibility is altered on the inactive X with aging and explore its potential escape from XCI.

The scope of our study is confined to two age groups, potentially missing any non-linear changes in chromatin accessibility with age. Still, reports on the transcriptomic changes in female microglia suggest a constituent trajectories across the aging process, implying a linear epigenetic regulation ^11,12^. Further comparisons of these data to other findings including DNA modifications, microglia enhancer maps, and sorting of microglia by reactivational phenotype will increase the depth of interpretation of results. Nevertheless, the analysis presented offers valuable insights into the mechanisms driving age-related microglia epigenetic changes in female mice.

In conclusion, these data present evidence of specific patterns with which female microglia respond to aging through shifts in chromatin accessibility. We report that alterations in chromatin accessibility at specific genomic sites involve both gains and losses, contrary to a global gain in accessibility. Still, the majority of changes in chromatin accessibility involve gaining accessibility rather than losing accessibility. Further, changes in accessibility are regulated by genomic region, which may affect the functional phenotype of female microglia with aging. Lastly, our data suggests that all chromosomes, including chromosome X, undergo age-related changes in accessibility, with a chromosome non-specific mechanism for increased accessibility in aged microglia.

## 5 Future directions

Future studies will explore sex differences in chromatin accessibility with aging and determine the independent contributions of sex chromosomal and gonadal hormones to these effects. It is important to highlight that currently we cannot distinguish deleterious and compensatory changes with age. We may employ chromatin immunoprecipitation–sequencing (ChIP-seq) to examine histone modifications, which would allow for the identification of regions associated with poised and active chromatin to enhance our ability to infer regulatory and functional networks. Lastly, with advances in epigenome editing, future functional studies will be conducted that examine the direct role of modifying accessibility at specific sites on microglia function during aging or disease.

## Data availability

The data from this study has been submitted and is now available in the GEO (Gene Expression Omnibus) repository under the accession code GSE251723.

## Acknowledgements

The authors wish to extend their gratitude to the Clinical Genomics Center (OMRF) and the Center for Biomedical Data Sciences (OMRF) for their assistance and instrument usage. Additionally, our appreciation goes to Adeline Machalinski for her management of the mouse colony and genotyping, as well as Robyn Berent for administrative support and overseeing lab management.

## Contributions

VAA designed and performed experiments; acquired, analyzed, and interpreted data; and wrote the manuscript. SRO designed work and experiments. KBT, KS, WH analyzed data; and revised the manuscript. KDP assisted in experiments and data processing; WMF designed the work; designed experiments; analyzed and interpreted data; and substantively revised the manuscript.

## Supplementary Information

**Additional file S1:** BamStats of samples. The read counts for samples used in this study, before and after subsampling.

**Additional file S2:** All peaks identified. Contains all the location of all ATAC peaks identified in this study.

**Additional file S3:** Specific and Shared sites. Consensus peaks common between young and old samples, young-specific and old-specific regions.

**Additional file S4:** Consensus peaks and differential peaks. Consensus peaks after re-centering peaks and the differentially accessible regions (DARs).

**Additional file S5:** DARs of samples from Li et al. study.

**Additional file S6:** Closest genes to DARs (± 3kb from gene)

**Additional file S7 – S10:** Homer motif enrichment files. Contains the *de novo* and known enriched motifs from Homer analysis.

**Additional file S11:** Differentially expressed genes identified in the study by Ocañas et al.

**Additional file S12:** DARs of the X chromosome.

## Supplementary figure legends

**Figure S1:**
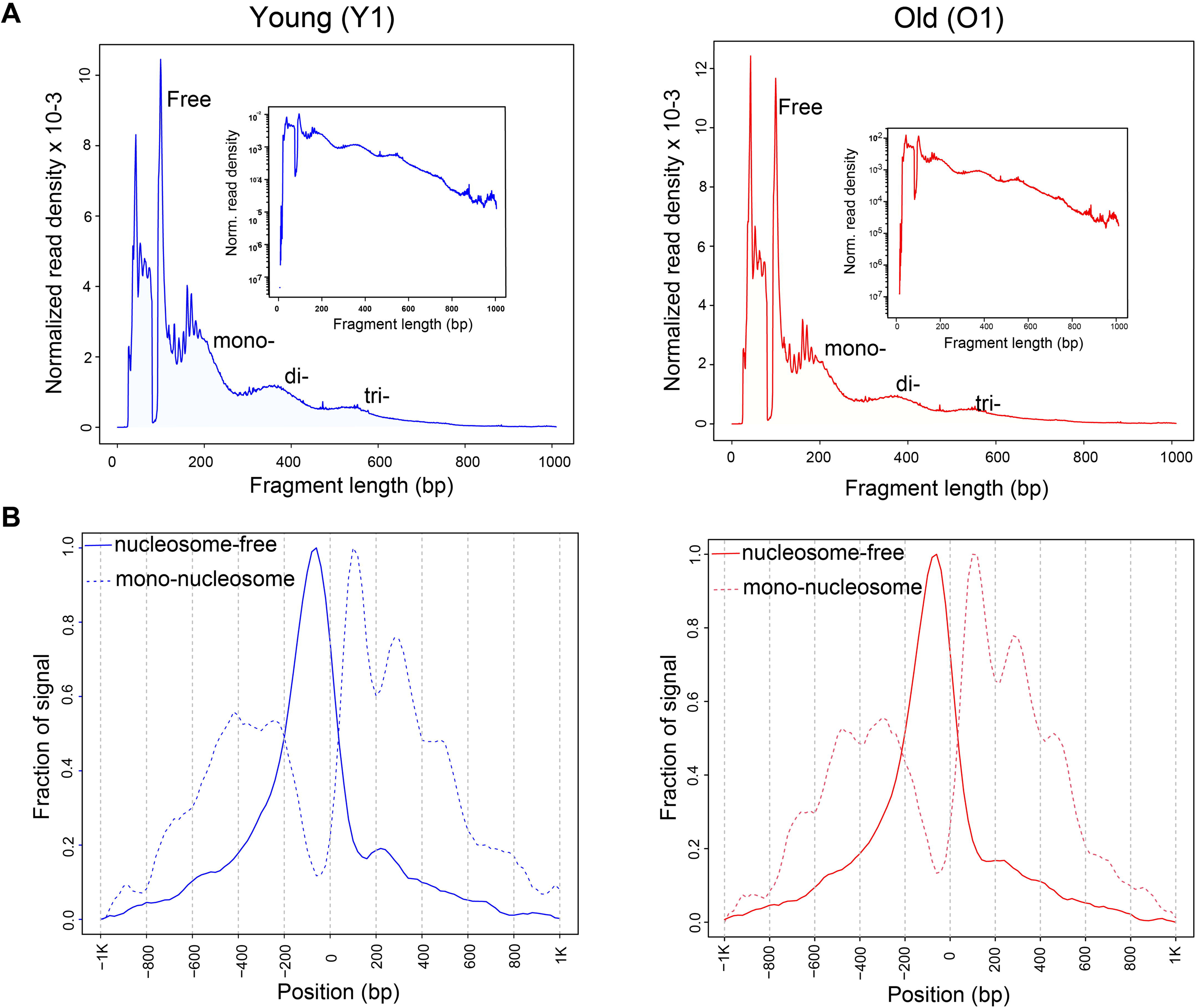
Fragment size distribution and nucleosome positioning. A) Distribution of fragment size was plotted for select young and old samples. We observe the highest number of fragments around 100 and 200 bp, suggesting an abundance of both nucleosome-free (Free) and mono-nucleosome-bound (mono-) fragments. Di-nucleosome (di-) and tri-nucleosome (tri-) bound fragments represented a smaller portion of fragment sizes. B) Nucleosome positioning and its relation to transcription start sites (TSS) was plotted for the same samples as above. Nucleosome-free (solid-line) fragments are more concentrated at the TSS, whereas mono-nucleosome (dashed-line) fragments are less prevalent at TSS but more abundant in neighboring regions.

**Figure S2:**
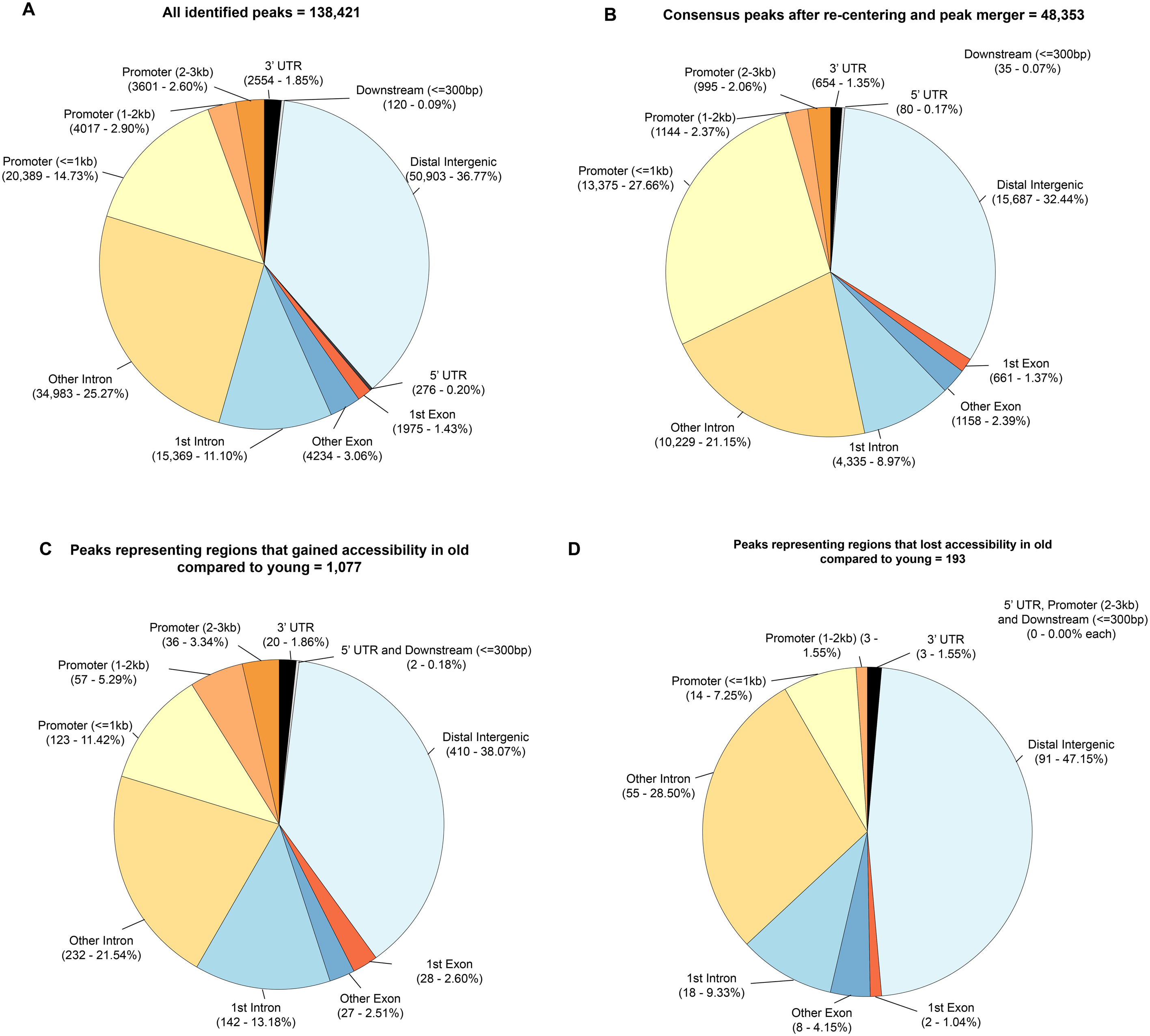
Genomic feature distributions of accessible chromatin regions. A) All identified peaks (138,421 peaks) B) Consensus peaks (48,353 peaks) C) Peaks that gained accessibility in old compared to young (1,077 peaks) and D) Peaks that lost accessibility in old compared to young (193 peaks).

**Figure S3:**
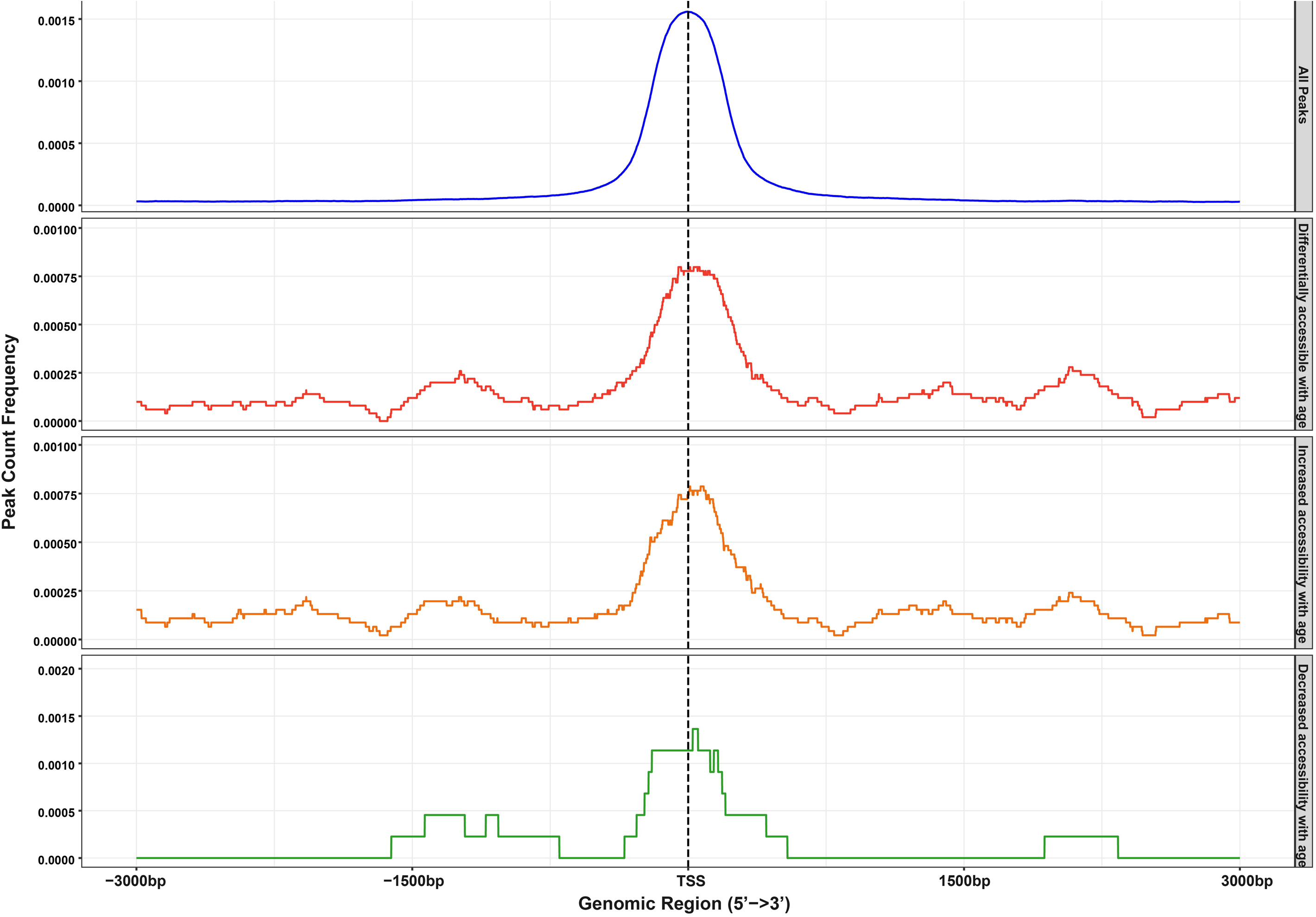
Peak density at the TSS. Whether examining all peaks or differentially accessible regions (DARs), a significant number of peaks exist at the transcription start site (TSS) of promoter regions. Nevertheless, DARs were under-represented at the proximal promoter was lower than expected in both regions that increased or decreased accessibility.

**Figure S4:**
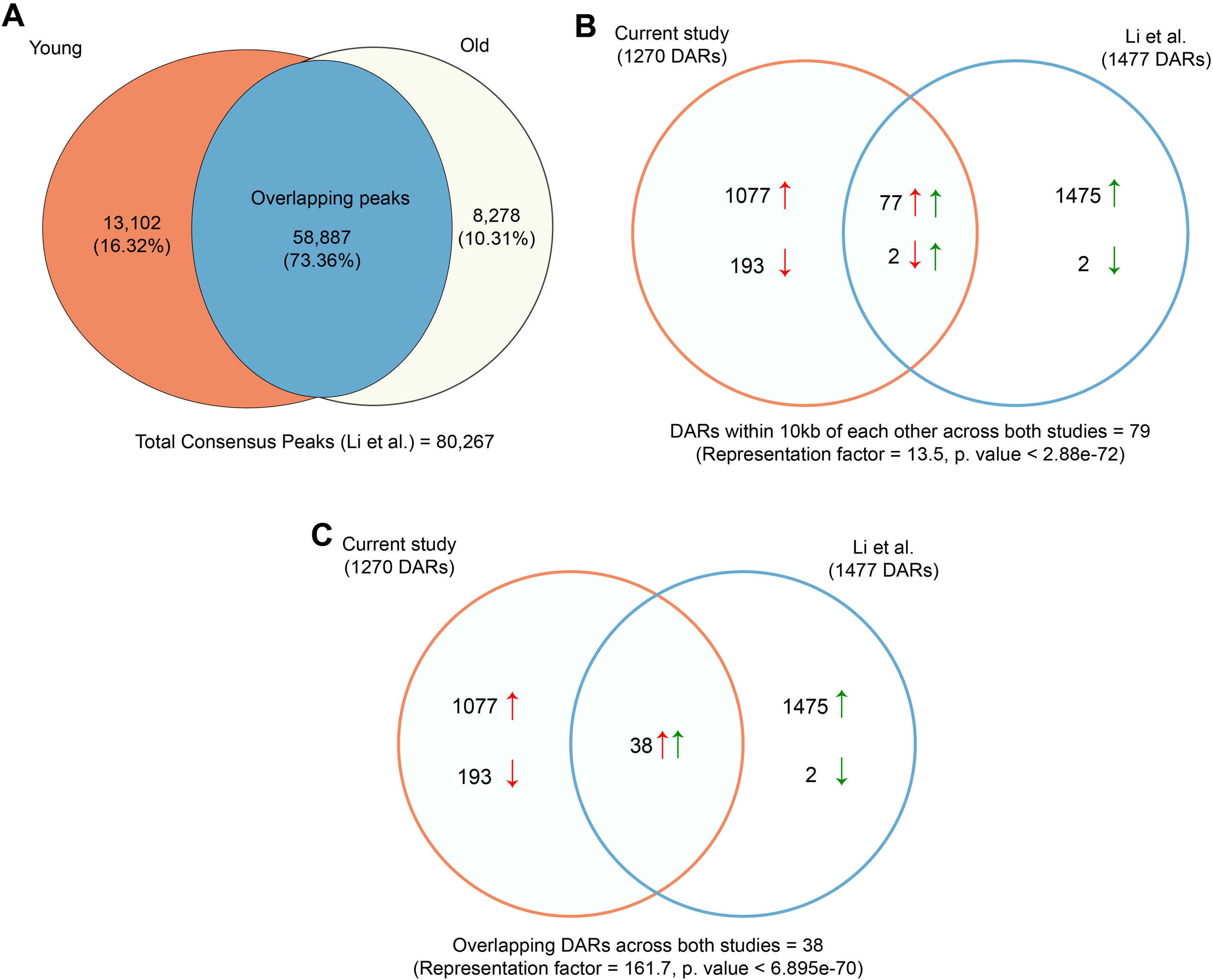
Comparative analysis with the study by Li et al. A) A set of consensus peaks totaling 80,267 were identified in microglia isolated from young (3 months) and old (24 months) PBS-treated mice. Within the consensus set, 58,887 peaks (representing 73.36%) were shared between the young and old mice, 13,102 peaks (16.32%) were exclusive to the young, and 8,278 peaks (10.31%) were exclusive to the old. Comparison of the statistical significance of changes in chromatin accessibility from Li et al and the present study that were within 10kb of each other (B) or overlapping (C). 79 DARs across the two studies were within 10kb of each other, 77 of which increased in accessibility in both studies. 2 DARs decreased accessibility in the present study but were found to be increasing accessibility in Li et al. ^11^

**Figure S5:**
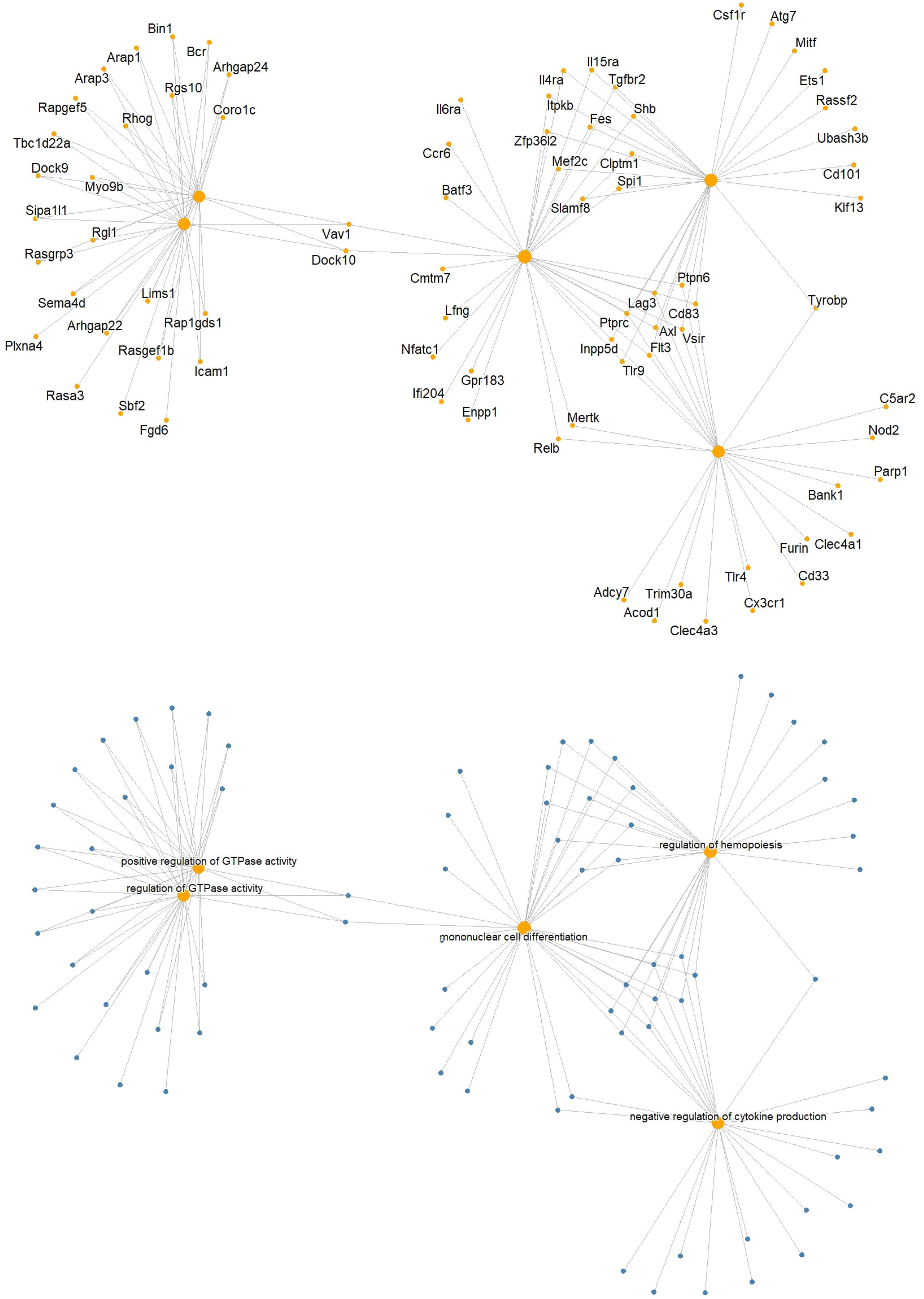
CNET plot of closest genes to regions that gained accessibility. CNET plot of genes closest (±3kb) to peaks increasing in accessibility in the top 5 pathways from the Gene Ontology analysis.

**Figure S6:**
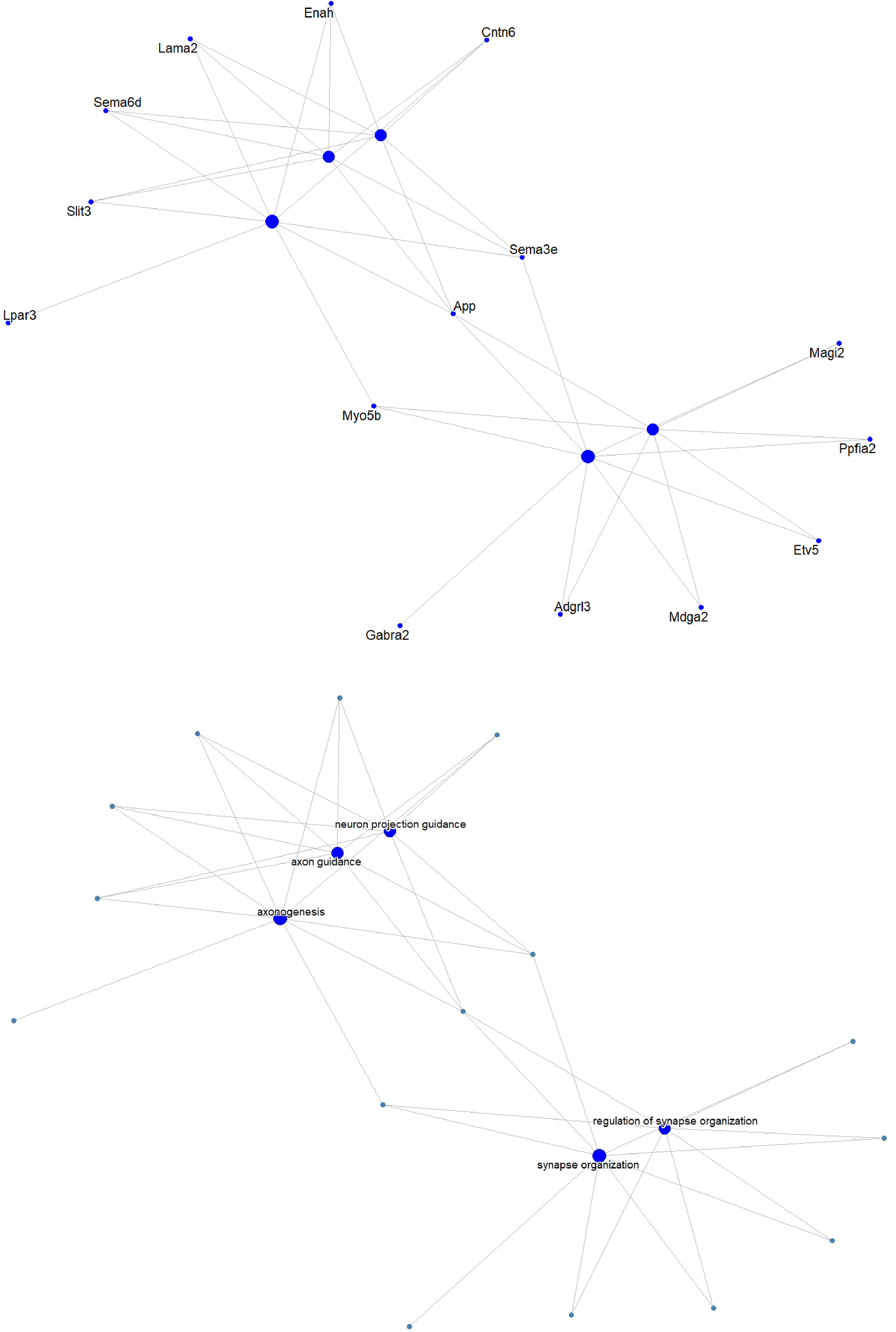
**CNET plot of closest genes to regions that lost accessibility**. CNET plot of genes closest (±3kb) to peaks decreasing in accessibility in the top 5 pathways from the Gene Ontology analysis.

