## Additional file S7-S10 for "Changes in microglia chromatin accessibility in aged female mice": Additional file S7_Homer_de novo motifs_gained.html

gainedHM// - Homer de novo Motif Results


### Homer *de novo* Motif Results (gainedHM//)

Known Motif Enrichment Results  
Gene Ontology Enrichment Results  
If Homer is having trouble matching a motif to a known motif, try copy/pasting the matrix file into
STAMP  
More information on motif finding results: HOMER
| Description of Results
| Tips
  
Total target sequences = 1077  
Total background sequences = 46753  
\* - possible false positive  

|  |  |  |  |  |  |  |  |  |
| --- | --- | --- | --- | --- | --- | --- | --- | --- |
| Rank | Motif | P-value | log P-pvalue | % of Targets | % of Background | STD(Bg STD) | Best Match/Details | Motif File |
| 1 | T C G A A T G C A C G T A C G T A G T C A G T C A G C T A G T C G A C T G C A T | 1e-279 | -6.434e+02 | 46.05% | 6.67% | 50.1bp (70.8bp) | PB0058.1\_Sfpi1\_1/Jaspar(0.972) More Information \| Similar Motifs Found | motif file (matrix) |
| 2 | G T C A G C T A T A C G A G C T C T A G C G T A C T G A C G T A | 1e-68 | -1.568e+02 | 22.28% | 6.07% | 53.9bp (68.5bp) | PU.1:IRF8(ETS:IRF)/pDC-Irf8-ChIP-Seq(GSE66899)/Homer(0.876) More Information \| Similar Motifs Found | motif file (matrix) |
| 3 | A T G C G C T A C A T G C G T A G T A C G T C A A T C G A C T G | 1e-28 | -6.511e+01 | 38.90% | 23.63% | 54.8bp (69.7bp) | Smad2(MAD)/ES-SMAD2-ChIP-Seq(GSE29422)/Homer(0.881) More Information \| Similar Motifs Found | motif file (matrix) |
| 4 | T G A C G A C T C G T A C G T A C T G A C G T A G C T A C A G T C T G A C T A G | 1e-26 | -6.036e+01 | 12.35% | 4.30% | 55.8bp (65.3bp) | MEF2A/MA0052.3/Jaspar(0.948) More Information \| Similar Motifs Found | motif file (matrix) |
| 5 | C G A T C G A T A T C G A G T C C A T G G A T C C G T A C G T A | 1e-25 | -5.948e+01 | 12.26% | 4.29% | 54.3bp (64.5bp) | CEBP(bZIP)/ThioMac-CEBPb-ChIP-Seq(GSE21512)/Homer(0.888) More Information \| Similar Motifs Found | motif file (matrix) |
| 6 | C A T G G A T C C G A T A C T G A G C T A C T G A T C G G A C T G C A T C G A T | 1e-25 | -5.775e+01 | 17.92% | 8.00% | 55.1bp (66.4bp) | RUNX-AML(Runt)/CD4+-PolII-ChIP-Seq(Barski\_et\_al.)/Homer(0.935) More Information \| Similar Motifs Found | motif file (matrix) |
| 7 | T G A C G C A T A T C G C G A T C T A G T A C G A C G T T A G C C G T A T A C G G T A C G C T A | 1e-21 | -4.866e+01 | 16.71% | 7.82% | 56.5bp (68.3bp) | NRL/MA0842.1/Jaspar(0.767) More Information \| Similar Motifs Found | motif file (matrix) |
| 8 | A G C T A C T G G T C A A T C G C G A T G T A C C T G A T A C G A T G C T A C G | 1e-20 | -4.752e+01 | 8.36% | 2.61% | 52.4bp (67.5bp) | Bach2(bZIP)/OCILy7-Bach2-ChIP-Seq(GSE44420)/Homer(0.908) More Information \| Similar Motifs Found | motif file (matrix) |
| 9 | C T A G T A C G C G A T G T A C G C T A A G T C T G C A G C A T A T C G G T C A | 1e-15 | -3.601e+01 | 5.29% | 1.45% | 52.6bp (64.1bp) | MITF(bHLH)/MastCells-MITF-ChIP-Seq(GSE48085)/Homer(0.896) More Information \| Similar Motifs Found | motif file (matrix) |
| 10 | T A G C C G T A T A C G C A G T G C A T A G C T G A T C T A C G | 1e-14 | -3.256e+01 | 13.37% | 6.74% | 55.1bp (68.2bp) | PB0034.1\_Irf4\_1/Jaspar(0.802) More Information \| Similar Motifs Found | motif file (matrix) |
| 11 | A T C G T A G C C T G A A T G C C G T A A T C G C G T A G T A C A G T C C G T A A T G C A C G T | 1e-13 | -3.106e+01 | 0.93% | 0.02% | 52.5bp (75.2bp) | PB0196.1\_Zbtb7b\_2/Jaspar(0.631) More Information \| Similar Motifs Found | motif file (matrix) |
| 12 | C T A G A C T G T A C G C G T A C T G A A C T G A C T G G T C A C T G A A C T G A G T C C G A T | 1e-13 | -3.077e+01 | 2.04% | 0.23% | 56.4bp (65.3bp) | EHF(ETS)/LoVo-EHF-ChIP-Seq(GSE49402)/Homer(0.643) More Information \| Similar Motifs Found | motif file (matrix) |
| 13 | T A G C G C T A A T G C C T A G A C T G G C T A C G T A A G T C G A C T C T G A | 1e-12 | -2.775e+01 | 4.64% | 1.42% | 53.1bp (67.8bp) | EHF/MA0598.2/Jaspar(0.645) More Information \| Similar Motifs Found | motif file (matrix) |
| 14 \* | C G T A C G T A C T A G T A G C C G A T C T A G C G T A T G C A T G A C T G A C C T G A A C T G | 1e-10 | -2.422e+01 | 2.14% | 0.36% | 48.0bp (83.7bp) | MafK(bZIP)/C2C12-MafK-ChIP-Seq(GSE36030)/Homer(0.614) More Information \| Similar Motifs Found | motif file (matrix) |
| 15 \* | A T C G G T A C T A G C C T A G A T G C G T C A C G T A A T C G A C T G A G T C | 1e-10 | -2.348e+01 | 1.95% | 0.31% | 54.4bp (69.9bp) | TFAP2B/MA0811.1/Jaspar(0.725) More Information \| Similar Motifs Found | motif file (matrix) |
| 16 \* | T G A C C G T A A T G C G A T C A G T C C G T A A C T G A C G T A C T G A C T G A C T G A C G T | 1e-8 | -2.022e+01 | 0.84% | 0.04% | 51.6bp (50.3bp) | ZKSCAN1(Zf)/HepG2-ZKSCAN1-ChIP-Seq(Encode)/Homer(0.707) More Information \| Similar Motifs Found | motif file (matrix) |
| 17 \* | A C G T A C T G A C T G T C A G G T A C A T C G T C A G T G C A A C G T A C T G | 1e-8 | -1.967e+01 | 2.14% | 0.46% | 57.4bp (60.1bp) | PB0143.1\_Klf7\_2/Jaspar(0.765) More Information \| Similar Motifs Found | motif file (matrix) |
| 18 \* | C A G T G T A C A C G T C T G A A C T G G T C A G A T C T A C G | 1e-6 | -1.469e+01 | 4.55% | 2.06% | 60.5bp (65.1bp) | SMAD3/MA0795.1/Jaspar(0.883) More Information \| Similar Motifs Found | motif file (matrix) |
| 19 \* | A C T G G T A C A C G T A C G T A G C T A C T G A G T C A C G T | 1e-6 | -1.398e+01 | 5.20% | 2.56% | 49.3bp (69.9bp) | SOX10/MA0442.2/Jaspar(0.701) More Information \| Similar Motifs Found | motif file (matrix) |
| 20 \* | C A T G G T C A G C A T A T C G A G C T G A T C G A C T A C T G A T G C A T C G | 1e-1 | -3.495e+00 | 0.28% | 0.06% | 43.7bp (61.4bp) | Smad4(MAD)/ESC-SMAD4-ChIP-Seq(GSE29422)/Homer(0.649) More Information \| Similar Motifs Found | motif file (matrix) |
