## Additional file S7-S10 for "Changes in microglia chromatin accessibility in aged female mice": Additional file S8_Homer_de novo motifs_loss.html

lostHM// - Homer de novo Motif Results


### Homer *de novo* Motif Results (lostHM//)

Known Motif Enrichment Results  
Gene Ontology Enrichment Results  
If Homer is having trouble matching a motif to a known motif, try copy/pasting the matrix file into
STAMP  
More information on motif finding results: HOMER
| Description of Results
| Tips
  
Total target sequences = 193  
Total background sequences = 47228  
\* - possible false positive  

|  |  |  |  |  |  |  |  |  |
| --- | --- | --- | --- | --- | --- | --- | --- | --- |
| Rank | Motif | P-value | log P-pvalue | % of Targets | % of Background | STD(Bg STD) | Best Match/Details | Motif File |
| 1 | G T A C A G T C C T G A A C G T T G A C A C G T A T C G A T C G A C G T A T C G | 1e-39 | -9.031e+01 | 34.72% | 4.60% | 50.5bp (64.4bp) | NEUROD1/MA1109.1/Jaspar(0.946) More Information \| Similar Motifs Found | motif file (matrix) |
| 2 | A G T C A G T C C T G A A G C T C A T G C T A G T A G C C G T A C T G A A G T C | 1e-14 | -3.355e+01 | 12.44% | 1.48% | 50.9bp (66.0bp) | Rfx1/MA0509.1/Jaspar(0.854) More Information \| Similar Motifs Found | motif file (matrix) |
| 3 | C G A T G A C T T G C A C G A T A T C G G A T C G T C A C G T A A G T C G T A C | 1e-13 | -3.164e+01 | 25.91% | 7.71% | 55.5bp (64.8bp) | HLF(bZIP)/HSC-HLF.Flag-ChIP-Seq(GSE69817)/Homer(0.869) More Information \| Similar Motifs Found | motif file (matrix) |
| 4 | C G T A T G C A C G T A C G T A A C T G C T G A G T C A A G C T G A T C A G T C C G T A C G T A | 1e-12 | -2.779e+01 | 8.81% | 0.82% | 53.5bp (64.7bp) | PB0037.1\_Isgf3g\_1/Jaspar(0.633) More Information \| Similar Motifs Found | motif file (matrix) |
| 5 \* | T G A C G T A C A G T C T C G A A G T C A C G T G A T C A C T G | 1e-11 | -2.711e+01 | 19.17% | 4.96% | 53.4bp (63.8bp) | PB0114.1\_Egr1\_2/Jaspar(0.859) More Information \| Similar Motifs Found | motif file (matrix) |
| 6 \* | C G T A C A T G A C T G C T G A A C T G C T A G A G T C C T A G A G C T C T A G | 1e-10 | -2.457e+01 | 5.70% | 0.29% | 53.9bp (64.7bp) | KLF14/MA0740.1/Jaspar(0.776) More Information \| Similar Motifs Found | motif file (matrix) |
| 7 \* | A G T C A G C T C A G T A T C G C G T A C G T A C T G A A C T G G A T C C T G A | 1e-10 | -2.411e+01 | 8.81% | 1.04% | 54.6bp (71.8bp) | BCL6B/MA0731.1/Jaspar(0.714) More Information \| Similar Motifs Found | motif file (matrix) |
| 8 \* | C G T A C T A G C T G A A G T C C G A T A C G T A C G T A G T C A C G T A C T G | 1e-10 | -2.400e+01 | 8.29% | 0.90% | 55.1bp (66.4bp) | ZNF768(Zf)/Rajj-ZNF768-ChIP-Seq(GSE111879)/Homer(0.681) More Information \| Similar Motifs Found | motif file (matrix) |
| 9 \* | C G T A A G C T A G C T G C A T C G T A G T C A C T G A A T C G C G T A C G T A G T A C C G T A | 1e-9 | -2.158e+01 | 10.36% | 1.78% | 54.7bp (64.3bp) | PB0123.1\_Foxl1\_2/Jaspar(0.697) More Information \| Similar Motifs Found | motif file (matrix) |
| 10 \* | C T A G A G T C A C G T A G C T C G A T C T A G A G T C C G A T A T C G T G C A | 1e-9 | -2.114e+01 | 7.77% | 0.94% | 54.9bp (71.9bp) | NRL/MA0842.1/Jaspar(0.712) More Information \| Similar Motifs Found | motif file (matrix) |
| 11 \* | A C T G A G T C C A G T C G T A A C G T C A G T A C G T C G T A A C T G C G A T | 1e-9 | -2.111e+01 | 7.77% | 0.94% | 48.5bp (60.7bp) | Ronin(THAP)/ES-Thap11-ChIP-Seq(GSE51522)/Homer(0.610) More Information \| Similar Motifs Found | motif file (matrix) |
| 12 \* | A T G C A T C G A G T C A C T G A G C T A C G T A C G T G A T C | 1e-8 | -2.047e+01 | 10.36% | 1.91% | 58.9bp (76.4bp) | Hes2/MA0616.1/Jaspar(0.658) More Information \| Similar Motifs Found | motif file (matrix) |
| 13 \* | C T G A C T A G T A C G T C G A C T G A C T G A C G T A C G T A C T G A C G T A C G T A A G C T | 1e-8 | -2.006e+01 | 9.33% | 1.55% | 55.4bp (71.1bp) | ZNF384/MA1125.1/Jaspar(0.820) More Information \| Similar Motifs Found | motif file (matrix) |
| 14 \* | T C A G G T C A C G A T A C G T G T A C C G T A A C G T A T G C A G T C G A C T | 1e-8 | -1.969e+01 | 8.29% | 1.22% | 51.7bp (68.4bp) | Fra2(bZIP)/Striatum-Fra2-ChIP-Seq(GSE43429)/Homer(0.701) More Information \| Similar Motifs Found | motif file (matrix) |
| 15 \* | T G C A C T G A G C A T A C G T G A C T C G T A A C T G A G T C | 1e-8 | -1.866e+01 | 27.98% | 12.54% | 52.9bp (71.9bp) | PB0145.1\_Mafb\_2/Jaspar(0.698) More Information \| Similar Motifs Found | motif file (matrix) |
| 16 \* | A C G T A G T C A G T C A G T C A G T C A C T G A C G T A C T G A G T C C G T A | 1e-7 | -1.623e+01 | 1.55% | 0.01% | 36.0bp (45.7bp) | PB0111.1\_Bhlhb2\_2/Jaspar(0.705) More Information \| Similar Motifs Found | motif file (matrix) |
| 17 \* | A C G T A G T C A C G T T G C A A C T G A G C T A G T C C T G A | 1e-6 | -1.524e+01 | 9.84% | 2.38% | 52.5bp (65.8bp) | SMAD3/MA0795.1/Jaspar(0.725) More Information \| Similar Motifs Found | motif file (matrix) |
| 18 \* | A C G T G A T C C G A T T A C G G A C T T A G C C G T A T C A G G T C A A C G T G T C A T G C A | 1e-6 | -1.478e+01 | 13.99% | 4.65% | 58.9bp (64.6bp) | MEIS2/MA0774.1/Jaspar(0.706) More Information \| Similar Motifs Found | motif file (matrix) |
| 19 \* | A G T C A C G T A C T G A C G T A C T G C T A G A C G T A C G T C T A G A G T C A C G T C G T A | 1e-5 | -1.170e+01 | 1.04% | 0.00% | 37.9bp (42.4bp) | RUNX-AML(Runt)/CD4+-PolII-ChIP-Seq(Barski\_et\_al.)/Homer(0.793) More Information \| Similar Motifs Found | motif file (matrix) |
