## Additional file S7-S10 for "Changes in microglia chromatin accessibility in aged female mice": Additional file S9_Homer_known motifs_gained.html

gainedHM/ - Homer Known Motif Enrichment Results


### Homer Known Motif Enrichment Results (gainedHM/)

Homer *de novo* Motif Results  
Gene Ontology Enrichment Results  
Known Motif Enrichment Results (txt file)  
Total Target Sequences = 1077, Total Background Sequences = 46752

|  |  |  |  |  |  |  |  |  |  |  |  |
| --- | --- | --- | --- | --- | --- | --- | --- | --- | --- | --- | --- |
| Rank | Motif | Name | P-value | log P-pvalue | q-value (Benjamini) | # Target Sequences with Motif | % of Targets Sequences with Motif | # Background Sequences with Motif | % of Background Sequences with Motif | Motif File | SVG |
| 1 | C G T A T A C G T C G A A C T G A C T G C G T A C G T A T A C G A G C T T A C G | PU.1(ETS)/ThioMac-PU.1-ChIP-Seq(GSE21512)/Homer | 1e-213 | -4.921e+02 | 0.0000 | 419.0 | 38.90% | 2889.3 | 6.18% | motif file (matrix) | svg |
| 2 | G C T A A G T C T A C G T G C A A T C G T C A G G C T A T C G A T C A G A G C T | ELF5(ETS)/T47D-ELF5-ChIP-Seq(GSE30407)/Homer | 1e-177 | -4.089e+02 | 0.0000 | 430.0 | 39.93% | 3842.2 | 8.22% | motif file (matrix) | svg |
| 3 | C T G A A G T C C G A T A G C T A T G C G T A C A C G T A T C G C A G T G C A T | Elf4(ETS)/BMDM-Elf4-ChIP-Seq(GSE88699)/Homer | 1e-171 | -3.945e+02 | 0.0000 | 514.0 | 47.73% | 5945.7 | 12.72% | motif file (matrix) | svg |
| 4 | C G T A C T G A C G T A C T A G T C G A C T A G A C T G C G T A C G T A T A C G A G C T A T C G | SpiB(ETS)/OCILY3-SPIB-ChIP-Seq(GSE56857)/Homer | 1e-163 | -3.776e+02 | 0.0000 | 262.0 | 24.33% | 1246.1 | 2.67% | motif file (matrix) | svg |
| 5 | T C G A T A G C T G C A A C T G A C T G C G T A C G T A C T A G G A C T T A C G | ETS1(ETS)/Jurkat-ETS1-ChIP-Seq(GSE17954)/Homer | 1e-131 | -3.024e+02 | 0.0000 | 449.0 | 41.69% | 5636.2 | 12.06% | motif file (matrix) | svg |
| 6 | C G T A T A G C T A G C T G C A A C T G C T A G C G T A C G T A T C A G G A C T | EHF(ETS)/LoVo-EHF-ChIP-Seq(GSE49402)/Homer | 1e-129 | -2.978e+02 | 0.0000 | 503.0 | 46.70% | 7170.9 | 15.35% | motif file (matrix) | svg |
| 7 | A T G C A G T C C T G A A G T C C G A T A C G T A G T C A G T C A C G T A T C G G A C T A C G T | Etv2(ETS)/ES-ER71-ChIP-Seq(GSE59402)/Homer | 1e-126 | -2.912e+02 | 0.0000 | 420.0 | 39.00% | 5075.8 | 10.86% | motif file (matrix) | svg |
| 8 | A G T C C T G A A G T C C G A T C A G T G A T C A T G C A C T G A T C G G A C T | Fli1(ETS)/CD8-FLI-ChIP-Seq(GSE20898)/Homer | 1e-125 | -2.897e+02 | 0.0000 | 446.0 | 41.41% | 5759.0 | 12.32% | motif file (matrix) | svg |
| 9 | C T A G C T A G C G T A C G T A T A C G C G A T C T A G C T G A C T G A C G T A T A C G G A C T | PU.1:IRF8(ETS:IRF)/pDC-Irf8-ChIP-Seq(GSE66899)/Homer | 1e-125 | -2.892e+02 | 0.0000 | 207.0 | 19.22% | 1009.1 | 2.16% | motif file (matrix) | svg |
| 10 | T C G A C T G A T A G C T G A C T C A G T C A G C G T A C G T A T C A G A G C T | ETV1(ETS)/GIST48-ETV1-ChIP-Seq(GSE22441)/Homer | 1e-125 | -2.885e+02 | 0.0000 | 517.0 | 48.00% | 7745.8 | 16.58% | motif file (matrix) | svg |
| 11 | C G T A T G A C T A G C T G C A A C T G A C T G C G T A C G T A T C A G G A C T | ELF3(ETS)/PDAC-ELF3-ChIP-Seq(GSE64557)/Homer | 1e-121 | -2.794e+02 | 0.0000 | 361.0 | 33.52% | 3849.1 | 8.24% | motif file (matrix) | svg |
| 12 | T C A G T C A G G C T A C G T A T A C G G A C T T C A G T C G A C T G A C G T A T A C G G A C T | IRF8(IRF)/BMDM-IRF8-ChIP-Seq(GSE77884)/Homer | 1e-121 | -2.791e+02 | 0.0000 | 245.0 | 22.75% | 1619.9 | 3.47% | motif file (matrix) | svg |
| 13 | T C G A T A G C G T C A A C T G A C T G C G T A C G T A C T A G A G C T T C A G | ERG(ETS)/VCaP-ERG-ChIP-Seq(GSE14097)/Homer | 1e-114 | -2.646e+02 | 0.0000 | 539.0 | 50.05% | 8897.7 | 19.04% | motif file (matrix) | svg |
| 14 | T C G A T C G A T A G C G T A C T C A G T A C G C G T A C G T A T C A G A G C T | GABPA(ETS)/Jurkat-GABPa-ChIP-Seq(GSE17954)/Homer | 1e-114 | -2.640e+02 | 0.0000 | 398.0 | 36.95% | 4943.3 | 10.58% | motif file (matrix) | svg |
| 15 | T G C A C T G A A T G C G T C A A C T G A C T G C G T A C G T A C T A G A G C T | Ets1-distal(ETS)/CD4+-PolII-ChIP-Seq(Barski\_et\_al.)/Homer | 1e-87 | -2.017e+02 | 0.0000 | 208.0 | 19.31% | 1648.3 | 3.53% | motif file (matrix) | svg |
| 16 | C T G A T A G C T G A C T C A G C T A G G T C A C G T A T C A G A G C T T C A G | ETV4(ETS)/HepG2-ETV4-ChIP-Seq(ENCODE)/Homer | 1e-86 | -2.002e+02 | 0.0000 | 394.0 | 36.58% | 5971.0 | 12.78% | motif file (matrix) | svg |
| 17 | T C G A A G C T A C G T A C G T A G T C A G T C A C G T A T C G G A C T A T C G | EWS:ERG-fusion(ETS)/CADO\_ES1-EWS:ERG-ChIP-Seq(SRA014231)/Homer | 1e-84 | -1.944e+02 | 0.0000 | 296.0 | 27.48% | 3554.6 | 7.61% | motif file (matrix) | svg |
| 18 | T G A C C T A G T C A G G T C A C G T A T C A G C G A T T C A G T C G A T G C A C T G A T A G C | PU.1-IRF(ETS:IRF)/Bcell-PU.1-ChIP-Seq(GSE21512)/Homer | 1e-83 | -1.928e+02 | 0.0000 | 462.0 | 42.90% | 8142.2 | 17.42% | motif file (matrix) | svg |
| 19 | C T G A T A C G G C A T A G C T A G C T A G T C T C G A A C T G C A G T A G C T A G C T G A T C | IRF3(IRF)/BMDM-Irf3-ChIP-Seq(GSE67343)/Homer | 1e-73 | -1.704e+02 | 0.0000 | 178.0 | 16.53% | 1417.8 | 3.03% | motif file (matrix) | svg |
| 20 | C T G A T G C A T A G C T G A C T A C G T C A G C T G A G C T A T C A G G A C T | ELF1(ETS)/Jurkat-ELF1-ChIP-Seq(SRA014231)/Homer | 1e-56 | -1.294e+02 | 0.0000 | 208.0 | 19.31% | 2533.7 | 5.42% | motif file (matrix) | svg |
| 21 | T G C A T C G A T A G C G T A C T C A G C T A G G T C A G C T A T C A G G A C T | ETS(ETS)/Promoter/Homer | 1e-49 | -1.135e+02 | 0.0000 | 152.0 | 14.11% | 1552.1 | 3.32% | motif file (matrix) | svg |
| 22 | T G C A C T G A A G T C G T C A A C T G A C T G C G T A C G T A C T G A A G C T | EWS:FLI1-fusion(ETS)/SK\_N\_MC-EWS:FLI1-ChIP-Seq(SRA014231)/Homer | 1e-44 | -1.017e+02 | 0.0000 | 212.0 | 19.68% | 3137.6 | 6.71% | motif file (matrix) | svg |
| 23 | T C A G C T G A C G T A C G T A T A C G G C A T C T A G C T G A C G T A C G T A T A C G G A C T | IRF1(IRF)/PBMC-IRF1-ChIP-Seq(GSE43036)/Homer | 1e-40 | -9.332e+01 | 0.0000 | 92.0 | 8.54% | 671.0 | 1.44% | motif file (matrix) | svg |
| 24 | G A T C T C G A A G T C C G A T C G A T A G T C A T G C A C T G A T C G G A C T | Elk1(ETS)/Hela-Elk1-ChIP-Seq(GSE31477)/Homer | 1e-34 | -7.934e+01 | 0.0000 | 172.0 | 15.97% | 2587.5 | 5.54% | motif file (matrix) | svg |
| 25 | C T G A A T G C C G T A A C G T A G T C A G T C A C G T A C T G A T C G G C A T | SPDEF(ETS)/VCaP-SPDEF-ChIP-Seq(SRA014231)/Homer | 1e-32 | -7.573e+01 | 0.0000 | 263.0 | 24.42% | 5271.6 | 11.28% | motif file (matrix) | svg |
| 26 | G A T C C T G A A G T C C G A T C G A T G A T C A G T C A C T G A T C G A G C T | Elk4(ETS)/Hela-Elk4-ChIP-Seq(GSE31477)/Homer | 1e-32 | -7.472e+01 | 0.0000 | 161.0 | 14.95% | 2407.0 | 5.15% | motif file (matrix) | svg |
| 27 | C T A G T C G A C T G A C G T A T A C G G A C T T C A G T C G A G T C A T G C A T A C G A G C T | IRF2(IRF)/Erythroblas-IRF2-ChIP-Seq(GSE36985)/Homer | 1e-27 | -6.379e+01 | 0.0000 | 65.0 | 6.04% | 494.4 | 1.06% | motif file (matrix) | svg |
| 28 | T C G A T A G C G T C A A C T G C T A G C G T A C G A T A C T G A C G T A C T G A C T G A C G T | ETS:RUNX(ETS,Runt)/Jurkat-RUNX1-ChIP-Seq(GSE17954)/Homer | 1e-23 | -5.320e+01 | 0.0000 | 52.0 | 4.83% | 378.9 | 0.81% | motif file (matrix) | svg |
| 29 | C A T G G T A C G A C T G C T A C G T A C G T A C G T A G C T A G A C T C T G A T C A G G T A C | Mef2c(MADS)/GM12878-Mef2c-ChIP-Seq(GSE32465)/Homer | 1e-22 | -5.186e+01 | 0.0000 | 83.0 | 7.71% | 985.1 | 2.11% | motif file (matrix) | svg |
| 30 | G A C T C T A G G A T C C A G T A C T G C T G A A T G C G C A T A T G C C T G A | MafA(bZIP)/Islet-MafA-ChIP-Seq(GSE30298)/Homer | 1e-20 | -4.693e+01 | 0.0000 | 207.0 | 19.22% | 4556.1 | 9.75% | motif file (matrix) | svg |
| 31 | C A T G A G T C G A C T C G T A C G A T G C A T G A C T G C A T C G A T C T A G C A T G T G A C | Mef2b(MADS)/HEK293-Mef2b.V5-ChIP-Seq(GSE67450)/Homer | 1e-20 | -4.643e+01 | 0.0000 | 132.0 | 12.26% | 2327.9 | 4.98% | motif file (matrix) | svg |
| 32 | G C T A T A G C A G C T A T C G G T C A C G T A G C T A A T G C G A T C C T G A | IRF4(IRF)/GM12878-IRF4-ChIP-Seq(GSE32465)/Homer | 1e-19 | -4.492e+01 | 0.0000 | 118.0 | 10.96% | 1986.3 | 4.25% | motif file (matrix) | svg |
| 33 | T G C A A G C T A C G T C T A G G A T C C T A G G A T C G T C A C T G A A G T C | CEBP(bZIP)/ThioMac-CEBPb-ChIP-Seq(GSE21512)/Homer | 1e-19 | -4.451e+01 | 0.0000 | 116.0 | 10.77% | 1943.4 | 4.16% | motif file (matrix) | svg |
| 34 | G T A C G A C T C G T A C T G A T C G A C G T A G C T A C A G T C T G A T A C G | Mef2a(MADS)/HL1-Mef2a.biotin-ChIP-Seq(GSE21529)/Homer | 1e-18 | -4.318e+01 | 0.0000 | 80.0 | 7.43% | 1067.3 | 2.28% | motif file (matrix) | svg |
| 35 | T A G C G C T A T C G A C T G A A G T C A G T C C T G A A G T C C G T A C T A G | RUNX(Runt)/HPC7-Runx1-ChIP-Seq(GSE22178)/Homer | 1e-17 | -4.133e+01 | 0.0000 | 166.0 | 15.41% | 3489.6 | 7.47% | motif file (matrix) | svg |
| 36 | A C T G G A T C G A C T A C T G A C G T C A T G A C T G A C G T A G C T C G A T | RUNX-AML(Runt)/CD4+-PolII-ChIP-Seq(Barski\_et\_al.)/Homer | 1e-16 | -3.841e+01 | 0.0000 | 161.0 | 14.95% | 3446.7 | 7.38% | motif file (matrix) | svg |
| 37 | T A C G T C A G A G C T A T G C C G T A A G T C T C A G A C G T A C T G T C G A | USF1(bHLH)/GM12878-Usf1-ChIP-Seq(GSE32465)/Homer | 1e-16 | -3.708e+01 | 0.0000 | 100.0 | 9.29% | 1709.5 | 3.66% | motif file (matrix) | svg |
| 38 | C T A G T C G A A C G T A C T G C G T A A T G C A C G T G T A C C G T A A G C T G A T C G T A C | Atf3(bZIP)/GBM-ATF3-ChIP-Seq(GSE33912)/Homer | 1e-15 | -3.662e+01 | 0.0000 | 133.0 | 12.35% | 2661.7 | 5.70% | motif file (matrix) | svg |
| 39 | A C T G C T A G T C G A C G A T C A T G G C T A A T C G C G A T G T A C G C T A A G C T G T A C | Fra1(bZIP)/BT549-Fra1-ChIP-Seq(GSE46166)/Homer | 1e-15 | -3.627e+01 | 0.0000 | 113.0 | 10.49% | 2092.5 | 4.48% | motif file (matrix) | svg |
| 40 | T A C G A T C G T A G C G A T C A C T G A C G T A G T C A C G T C T A G A T C G | Smad4(MAD)/ESC-SMAD4-ChIP-Seq(GSE29422)/Homer | 1e-15 | -3.560e+01 | 0.0000 | 322.0 | 29.90% | 9138.5 | 19.56% | motif file (matrix) | svg |
| 41 | A T G C G A C T A C T G C A G T G A T C A C G T T A C G T A C G | Smad2(MAD)/ES-SMAD2-ChIP-Seq(GSE29422)/Homer | 1e-15 | -3.546e+01 | 0.0000 | 321.0 | 29.81% | 9110.4 | 19.50% | motif file (matrix) | svg |
| 42 | G C T A C T G A T C G A A G T C A G T C C T G A A G T C G T C A C T G A T G C A | RUNX1(Runt)/Jurkat-RUNX1-ChIP-Seq(GSE29180)/Homer | 1e-14 | -3.316e+01 | 0.0000 | 208.0 | 19.31% | 5216.1 | 11.16% | motif file (matrix) | svg |
| 43 | C A G T T G C A A C G T A C T G C G T A A T C G C G A T T G A C C G T A A C G T | BATF(bZIP)/Th17-BATF-ChIP-Seq(GSE39756)/Homer | 1e-14 | -3.283e+01 | 0.0000 | 129.0 | 11.98% | 2677.7 | 5.73% | motif file (matrix) | svg |
| 44 | C G A T T A C G T G A C G A C T C A T G C G T A T A C G A C G T G T A C C T G A | Bach2(bZIP)/OCILy7-Bach2-ChIP-Seq(GSE44420)/Homer | 1e-14 | -3.280e+01 | 0.0000 | 64.0 | 5.94% | 891.3 | 1.91% | motif file (matrix) | svg |
| 45 | C A T G A G T C A G C T C G T A C G A T C G A T G C A T G C A T C G A T C T G A C A T G T G A C | Mef2d(MADS)/Retina-Mef2d-ChIP-Seq(GSE61391)/Homer | 1e-14 | -3.248e+01 | 0.0000 | 41.0 | 3.81% | 399.7 | 0.86% | motif file (matrix) | svg |
| 46 | C T A G T C G A G C A T C A T G G C T A T A G C C G A T G T A C C T G A A G C T | JunB(bZIP)/DendriticCells-Junb-ChIP-Seq(GSE36099)/Homer | 1e-13 | -3.162e+01 | 0.0000 | 111.0 | 10.31% | 2184.1 | 4.67% | motif file (matrix) | svg |
| 47 | C T A G T C G A C G A T A C T G C G T A T A C G A G C T T G A C G C T A A C G T G A T C T A G C | Fosl2(bZIP)/3T3L1-Fosl2-ChIP-Seq(GSE56872)/Homer | 1e-13 | -3.050e+01 | 0.0000 | 82.0 | 7.61% | 1402.4 | 3.00% | motif file (matrix) | svg |
| 48 | C T G A A T C G A G C T A G C T A C G T T A G C C T G A T A C G C G A T A C G T G A C T A G T C | ISRE(IRF)/ThioMac-LPS-Expression(GSE23622)/Homer | 1e-12 | -2.958e+01 | 0.0000 | 30.0 | 2.79% | 234.3 | 0.50% | motif file (matrix) | svg |
| 49 | C A T G C T A G T C G A A C G T A C T G C G T A T A G C C G A T T G A C C G T A A G C T G A T C | Fra2(bZIP)/Striatum-Fra2-ChIP-Seq(GSE43429)/Homer | 1e-12 | -2.881e+01 | 0.0000 | 98.0 | 9.10% | 1900.8 | 4.07% | motif file (matrix) | svg |
| 50 | T C G A A C G T C A T G G C T A T A G C C G A T G T A C G C T A A C G T A T G C | AP-1(bZIP)/ThioMac-PU.1-ChIP-Seq(GSE21512)/Homer | 1e-11 | -2.737e+01 | 0.0000 | 135.0 | 12.53% | 3085.3 | 6.60% | motif file (matrix) | svg |
| 51 | C T A G T C G A A C G T A C T G C G T A T A G C C G A T G T A C C G T A A G C T G A T C G T A C | Jun-AP1(bZIP)/K562-cJun-ChIP-Seq(GSE31477)/Homer | 1e-11 | -2.542e+01 | 0.0000 | 63.0 | 5.85% | 1032.3 | 2.21% | motif file (matrix) | svg |
| 52 | G T A C C G T A C G T A T A C G G C A T G T A C C G T A C A T G A G T C C G T A C G T A C G A T G C A T G C A T G A C T | MafF(bZIP)/HepG2-MafF-ChIP-Seq(GSE31477)/Homer | 1e-10 | -2.428e+01 | 0.0000 | 54.0 | 5.01% | 829.7 | 1.78% | motif file (matrix) | svg |
| 53 | T G A C G C T A T C G A T G C A A G T C A G T C C G T A A G T C C G T A C T G A G C T A G T A C | RUNX2(Runt)/PCa-RUNX2-ChIP-Seq(GSE33889)/Homer | 1e-10 | -2.417e+01 | 0.0000 | 162.0 | 15.04% | 4135.1 | 8.85% | motif file (matrix) | svg |
| 54 | T C G A T G A C G C A T A G C T C A G T G A T C G C T A G A T C G A C T A C G T G C A T A G T C | PRDM1(Zf)/Hela-PRDM1-ChIP-Seq(GSE31477)/Homer | 1e-9 | -2.256e+01 | 0.0000 | 122.0 | 11.33% | 2889.0 | 6.18% | motif file (matrix) | svg |
| 55 | A G C T G C A T A C T G A C G T A G T C A C G T C T A G T A C G | Smad3(MAD)/NPC-Smad3-ChIP-Seq(GSE36673)/Homer | 1e-9 | -2.125e+01 | 0.0000 | 476.0 | 44.20% | 16431.4 | 35.16% | motif file (matrix) | svg |
| 56 | T C A G A G C T A T G C C G T A A G C T T C A G C A G T A C T G C T G A A G T C | MITF(bHLH)/MastCells-MITF-ChIP-Seq(GSE48085)/Homer | 1e-9 | -2.124e+01 | 0.0000 | 152.0 | 14.11% | 3961.5 | 8.48% | motif file (matrix) | svg |
| 57 | T G C A A T G C A C G T A C G T A C G T A T G C C T A G A C G T A C G T A G C T G A T C A G C T | T1ISRE(IRF)/ThioMac-Ifnb-Expression/Homer | 1e-9 | -2.106e+01 | 0.0000 | 12.0 | 1.11% | 43.6 | 0.09% | motif file (matrix) | svg |
| 58 | C T G A T C A G G T A C G C T A A C T G T G A C G C A T C A T G | SCL(bHLH)/HPC7-Scl-ChIP-Seq(GSE13511)/Homer | 1e-8 | -2.037e+01 | 0.0000 | 657.0 | 61.00% | 24280.2 | 51.96% | motif file (matrix) | svg |
| 59 | T C A G T A G C G A C T C A T G C T G A A T C G G C A T G T A C C G T A A C T G T A G C T G C A | MafK(bZIP)/C2C12-MafK-ChIP-Seq(GSE36030)/Homer | 1e-8 | -2.036e+01 | 0.0000 | 71.0 | 6.59% | 1403.8 | 3.00% | motif file (matrix) | svg |
| 60 | T C A G G A C T C A G T C T G A A G C T C T A G G A C T T G C A C T G A A G T C | HLF(bZIP)/HSC-HLF.Flag-ChIP-Seq(GSE69817)/Homer | 1e-8 | -1.890e+01 | 0.0000 | 98.0 | 9.10% | 2290.4 | 4.90% | motif file (matrix) | svg |
| 61 | C T A G T C G A C G A T C T A G G C A T C A G T C T A G G A T C C G T A G T C A | CEBP:AP1(bZIP)/ThioMac-CEBPb-ChIP-Seq(GSE21512)/Homer | 1e-8 | -1.846e+01 | 0.0000 | 112.0 | 10.40% | 2768.0 | 5.92% | motif file (matrix) | svg |
| 62 | A G T C A C G T A C T G A G C T A C G T A C G T G T C A A G T C | Foxo1(Forkhead)/RAW-Foxo1-ChIP-Seq(Fan\_et\_al.)/Homer | 1e-7 | -1.836e+01 | 0.0000 | 296.0 | 27.48% | 9492.4 | 20.31% | motif file (matrix) | svg |
| 63 | C G T A G A C T C G A T A T C G G T A C G C A T C A T G C G T A T A C G G C A T G T A C C G T A C A T G A T G C G C T A C T A G G C A T G C A T G C A T G A C T | MafB(bZIP)/BMM-Mafb-ChIP-Seq(GSE75722)/Homer | 1e-7 | -1.758e+01 | 0.0000 | 94.0 | 8.73% | 2225.2 | 4.76% | motif file (matrix) | svg |
| 64 | T A C G T C G A C A G T A C T G G C T A A T G C C G A T G T A C C G T A A C T G T A G C C G T A | NF-E2(bZIP)/K562-NFE2-ChIP-Seq(GSE31477)/Homer | 1e-7 | -1.747e+01 | 0.0000 | 24.0 | 2.23% | 263.6 | 0.56% | motif file (matrix) | svg |
| 65 | T C A G A C G T A G T C T C G A A G T C T C A G G C A T C T A G C T A G A G C T | Usf2(bHLH)/C2C12-Usf2-ChIP-Seq(GSE36030)/Homer | 1e-7 | -1.621e+01 | 0.0000 | 56.0 | 5.20% | 1111.5 | 2.38% | motif file (matrix) | svg |
| 66 | A T G C G A C T A G C T A G C T A G T C G C T A C A G T C G A T G C T A A C G T A C T G G C T A T A G C G C A T T G A C | IRF:BATF(IRF:bZIP)/pDC-Irf8-ChIP-Seq(GSE66899)/Homer | 1e-7 | -1.621e+01 | 0.0000 | 30.0 | 2.79% | 417.6 | 0.89% | motif file (matrix) | svg |
| 67 | T C G A A C T G A C T G C G T A C G T A T C G A A G T C C T G A A T C G G T A C G C A T C A T G | ETS:E-box(ETS,bHLH)/HPC7-Scl-ChIP-Seq(GSE22178)/Homer | 1e-6 | -1.516e+01 | 0.0000 | 31.0 | 2.88% | 463.7 | 0.99% | motif file (matrix) | svg |
| 68 | G T C A T G C A G C T A A G T C C G T A A C T G T G A C G C A T T C A G C A G T | Ap4(bHLH)/AML-Tfap4-ChIP-Seq(GSE45738)/Homer | 1e-6 | -1.478e+01 | 0.0000 | 214.0 | 19.87% | 6685.6 | 14.31% | motif file (matrix) | svg |
| 69 | G T C A G C A T A C T G G T A C G A C T A C T G G C T A A T C G C A G T G T A C C G T A A G C T | Nrf2(bZIP)/Lymphoblast-Nrf2-ChIP-Seq(GSE37589)/Homer | 1e-6 | -1.417e+01 | 0.0000 | 19.0 | 1.76% | 208.8 | 0.45% | motif file (matrix) | svg |
| 70 | T C G A A C G T A C G T C T G A G A T C T C A G G A C T G T C A C G T A A G C T G T C A C T A G A G C T A C G T T C G A | NFIL3(bZIP)/HepG2-NFIL3-ChIP-Seq(Encode)/Homer | 1e-6 | -1.402e+01 | 0.0000 | 69.0 | 6.41% | 1595.2 | 3.41% | motif file (matrix) | svg |
| 71 | G A C T A G T C C G A T A C T G C T G A T G A C G T A C C G T A A T C G G C A T C T G A C T A G | Bcl11a(Zf)/HSPC-BCL11A-ChIP-Seq(GSE104676)/Homer | 1e-5 | -1.368e+01 | 0.0000 | 126.0 | 11.70% | 3541.8 | 7.58% | motif file (matrix) | svg |
| 72 | C T A G G C A T G A T C C G T A A G T C T C A G G A C T C T A G | CLOCK(bHLH)/Liver-Clock-ChIP-Seq(GSE39860)/Homer | 1e-5 | -1.277e+01 | 0.0000 | 84.0 | 7.80% | 2149.8 | 4.60% | motif file (matrix) | svg |
| 73 | T G C A A G C T C A T G C G T A A G C T A C T G G A T C G T C A C G T A A G C T | Atf4(bZIP)/MEF-Atf4-ChIP-Seq(GSE35681)/Homer | 1e-5 | -1.154e+01 | 0.0001 | 40.0 | 3.71% | 810.8 | 1.74% | motif file (matrix) | svg |
| 74 | C T A G T A C G G A C T T G C A T G C A C G A T T A C G C T G A T C G A C T G A | Hoxa10(Homeobox)/ChickenMSG-Hoxa10.Flag-ChIP-Seq(GSE86088)/Homer | 1e-4 | -1.136e+01 | 0.0001 | 78.0 | 7.24% | 2030.2 | 4.34% | motif file (matrix) | svg |
| 75 | C T A G C A G T T G A C C G T A G A T C T C A G G A C T C A T G | BMAL1(bHLH)/Liver-Bmal1-ChIP-Seq(GSE39860)/Homer | 1e-4 | -1.050e+01 | 0.0002 | 247.0 | 22.93% | 8421.1 | 18.02% | motif file (matrix) | svg |
| 76 | T C A G A G C T A T G C C G T A A G T C T C A G A C G T A T C G T C G A A G T C G A T C T G A C | TFE3(bHLH)/MEF-TFE3-ChIP-Seq(GSE75757)/Homer | 1e-4 | -1.044e+01 | 0.0002 | 17.0 | 1.58% | 225.8 | 0.48% | motif file (matrix) | svg |
| 77 | T C G A T A G C T G A C C T G A A G T C A C T G G A C T C A T G | c-Myc(bHLH)/LNCAP-cMyc-ChIP-Seq(Unpublished)/Homer | 1e-4 | -1.043e+01 | 0.0002 | 61.0 | 5.66% | 1516.9 | 3.25% | motif file (matrix) | svg |
| 78 | A T G C A T G C A T C G T A C G A G C T A G T C G C T A A G T C T C A G G A C T A C T G T C G A | E-box(bHLH)/Promoter/Homer | 1e-4 | -1.012e+01 | 0.0002 | 17.0 | 1.58% | 231.0 | 0.49% | motif file (matrix) | svg |
| 79 | C G A T C T A G C T G A A T G C C T G A T C G A C G T A C T G A T C G A T A G C A G T C C G T A A C T G T C G A A T G C | Hand2(bHLH)/Mesoderm-Hand2-ChIP-Seq(GSE61475)/Homer | 1e-4 | -9.946e+00 | 0.0003 | 93.0 | 8.64% | 2644.1 | 5.66% | motif file (matrix) | svg |
| 80 | C G T A C G T A C G T A G C A T G C A T T A C G G T A C G A C T A C T G C G T A A T C G A C G T G T A C C G T A A G C T | Bach1(bZIP)/K562-Bach1-ChIP-Seq(GSE31477)/Homer | 1e-4 | -9.527e+00 | 0.0004 | 16.0 | 1.49% | 219.8 | 0.47% | motif file (matrix) | svg |
| 81 | C A T G T G A C C G T A A G T C T A C G G C A T A C T G G T C A A T G C A G T C | bHLHE41(bHLH)/proB-Bhlhe41-ChIP-Seq(GSE93764)/Homer | 1e-4 | -9.518e+00 | 0.0004 | 141.0 | 13.09% | 4437.1 | 9.50% | motif file (matrix) | svg |
| 82 | A G T C G A C T C A G T A C T G C T A G T G A C G C T A A T G C G C A T A T C G C G A T A C T G G A T C G T A C G T C A C T G A | NF1(CTF)/LNCAP-NF1-ChIP-Seq(Unpublished)/Homer | 1e-3 | -9.191e+00 | 0.0005 | 67.0 | 6.22% | 1790.0 | 3.83% | motif file (matrix) | svg |
| 83 | C G A T T G C A G T A C C G T A A G T C C T A G G A C T C A T G | NPAS(bHLH)/Liver-NPAS-ChIP-Seq(GSE39860)/Homer | 1e-3 | -8.493e+00 | 0.0011 | 210.0 | 19.50% | 7217.5 | 15.45% | motif file (matrix) | svg |
| 84 | C A T G G T A C C G T A A G T C C T A G A C G T A C T G G T A C A G T C A G C T | bHLHE40(bHLH)/HepG2-BHLHE40-ChIP-Seq(GSE31477)/Homer | 1e-3 | -8.035e+00 | 0.0017 | 44.0 | 4.09% | 1087.6 | 2.33% | motif file (matrix) | svg |
| 85 | A G C T C A T G G C A T G A T C T G C A C T A G G A T C A C G T | Tgif2(Homeobox)/mES-Tgif2-ChIP-Seq(GSE55404)/Homer | 1e-3 | -8.004e+00 | 0.0017 | 427.0 | 39.65% | 16179.0 | 34.62% | motif file (matrix) | svg |
| 86 | T A C G T C G A T A G C A G T C C G T A A G T C C T A G G C A T A C T G A T C G | n-Myc(bHLH)/mES-nMyc-ChIP-Seq(GSE11431)/Homer | 1e-3 | -7.941e+00 | 0.0018 | 100.0 | 9.29% | 3062.1 | 6.55% | motif file (matrix) | svg |
| 87 | T C G A G C A T A C G T C T A G G T A C T C G A G C A T T G A C T C G A A C G T | Chop(bZIP)/MEF-Chop-ChIP-Seq(GSE35681)/Homer | 1e-3 | -7.897e+00 | 0.0018 | 29.0 | 2.69% | 620.9 | 1.33% | motif file (matrix) | svg |
| 88 | T A C G T C G A G A C T A C T G C T G A A G T C T C A G G A C T T G A C C T G A | Atf1(bZIP)/K562-ATF1-ChIP-Seq(GSE31477)/Homer | 1e-3 | -7.815e+00 | 0.0020 | 79.0 | 7.34% | 2309.5 | 4.94% | motif file (matrix) | svg |
| 89 | C T G A C T A G A T C G G C A T A C T G G T A C A T G C C G T A A C T G G C T A A G T C C G T A | Tbox:Smad(T-box,MAD)/ESCd5-Smad2\_3-ChIP-Seq(GSE29422)/Homer | 1e-3 | -7.717e+00 | 0.0021 | 39.0 | 3.62% | 941.1 | 2.01% | motif file (matrix) | svg |
| 90 | T C G A T C G A T A C G G A T C G T C A G T A C C G A T A G C T G T C A T G C A | Nkx3.1(Homeobox)/LNCaP-Nkx3.1-ChIP-Seq(GSE28264)/Homer | 1e-3 | -7.298e+00 | 0.0032 | 329.0 | 30.55% | 12218.6 | 26.15% | motif file (matrix) | svg |
| 91 | C G T A C A T G C A T G A C T G C T A G T C G A G C A T C G A T A G C T A G T C G A T C G T A C | NFkB-p65(RHD)/GM12787-p65-ChIP-Seq(GSE19485)/Homer | 1e-3 | -6.961e+00 | 0.0045 | 75.0 | 6.96% | 2235.9 | 4.78% | motif file (matrix) | svg |
| 92 | A C T G C A T G G C T A T C G A G C T A A G C T A G C T G T A C A G T C T G A C | NFkB-p65-Rel(RHD)/ThioMac-LPS-Expression(GSE23622)/Homer | 1e-2 | -6.675e+00 | 0.0059 | 14.0 | 1.30% | 232.9 | 0.50% | motif file (matrix) | svg |
| 93 | T C G A T G A C G T A C C G T A C A G T T G A C A C G T A C T G A G C T A G C T | NeuroG2(bHLH)/Fibroblast-NeuroG2-ChIP-Seq(GSE75910)/Homer | 1e-2 | -6.361e+00 | 0.0079 | 214.0 | 19.87% | 7685.1 | 16.45% | motif file (matrix) | svg |
| 94 | T C A G A C G T T C G A T A G C A G T C C G T A A C T G G T A C A C G T A C T G A T C G A G T C | Atoh1(bHLH)/Cerebellum-Atoh1-ChIP-Seq(GSE22111)/Homer | 1e-2 | -6.242e+00 | 0.0089 | 158.0 | 14.67% | 5473.5 | 11.71% | motif file (matrix) | svg |
| 95 | C T G A T C A G A G T C C G T A A T C G A T G C C G A T A C T G A G T C G A C T A T C G A G T C | MyoD(bHLH)/Myotube-MyoD-ChIP-Seq(GSE21614)/Homer | 1e-2 | -6.108e+00 | 0.0100 | 122.0 | 11.33% | 4087.5 | 8.75% | motif file (matrix) | svg |
| 96 | T C G A T G A C A G T C C G T A A G T C C T A G A C G T A C T G A C T G A G C T A G T C G C A T | Max(bHLH)/K562-Max-ChIP-Seq(GSE31477)/Homer | 1e-2 | -5.983e+00 | 0.0112 | 94.0 | 8.73% | 3032.8 | 6.49% | motif file (matrix) | svg |
| 97 | C T A G T C G A T G A C A G T C C G T A A C T G G T A C A C G T A C T G A C T G | BHLHA15(bHLH)/NIH3T3-BHLHB8.HA-ChIP-Seq(GSE119782)/Homer | 1e-2 | -5.893e+00 | 0.0122 | 197.0 | 18.29% | 7077.6 | 15.15% | motif file (matrix) | svg |
| 98 | C T G A T C A G G T A C T G C A A G T C C G T A A G T C A C T G A C G T A C T G | MNT(bHLH)/HepG2-MNT-ChIP-Seq(Encode)/Homer | 1e-2 | -5.775e+00 | 0.0136 | 135.0 | 12.53% | 4635.9 | 9.92% | motif file (matrix) | svg |
| 99 | T A G C C T A G T C G A G A C T A C T G C T G A A G T C T C A G G C A T T G A C C T G A A G C T | Atf7(bZIP)/3T3L1-Atf7-ChIP-Seq(GSE56872)/Homer | 1e-2 | -5.605e+00 | 0.0159 | 56.0 | 5.20% | 1662.2 | 3.56% | motif file (matrix) | svg |
| 100 | C T A G A C G T A G T C C G T A A C T G A G T C G C A T A C T G G C A T A G T C G A C T G A T C G C A T A G T C A G C T | ZNF317(Zf)/HEK293-ZNF317.GFP-ChIP-Seq(GSE58341)/Homer | 1e-2 | -5.562e+00 | 0.0164 | 20.0 | 1.86% | 440.6 | 0.94% | motif file (matrix) | svg |
| 101 | T C A G T C A G T A G C A G T C C T G A A G T C C T A G A C G T A C T G A T C G | c-Myc(bHLH)/mES-cMyc-ChIP-Seq(GSE11431)/Homer | 1e-2 | -5.373e+00 | 0.0197 | 75.0 | 6.96% | 2382.6 | 5.10% | motif file (matrix) | svg |
| 102 | C G T A C T G A C T A G C G T A C G T A A G T C C G T A C A G T G C A T G T C A C G A T A C T G A C G T G C A T G A T C | PGR(NR)/EndoStromal-PGR-ChIP-Seq(GSE69539)/Homer | 1e-2 | -5.323e+00 | 0.0205 | 36.0 | 3.34% | 977.8 | 2.09% | motif file (matrix) | svg |
| 103 | A C T G C T A G G A T C C G T A C T A G A G T C T G C A G A C T C G T A A C G T T C A G A G T C A C G T C T G A A G T C A G T C G A T C C G T A T C A G T A C G | EBNA1(EBV-virus)/Raji-EBNA1-ChIP-Seq(GSE30709)/Homer | 1e-2 | -5.214e+00 | 0.0226 | 6.0 | 0.56% | 68.2 | 0.15% | motif file (matrix) | svg |
| 104 | T A C G T G C A A G T C C G T A A C G T T G A C A C G T A C T G A C T G G C A T | TCF4(bHLH)/SHSY5Y-TCF4-ChIP-Seq(GSE96915)/Homer | 1e-2 | -5.149e+00 | 0.0239 | 212.0 | 19.68% | 7811.2 | 16.72% | motif file (matrix) | svg |
| 105 | T C G A A G T C C G T A A T C G A T G C C G A T A C T G A G T C A G C T A C T G | Tcf12(bHLH)/GM12878-Tcf12-ChIP-Seq(GSE32465)/Homer | 1e-2 | -5.142e+00 | 0.0239 | 146.0 | 13.56% | 5159.6 | 11.04% | motif file (matrix) | svg |
| 106 | T C A G A T C G G A C T A C T G G A C T C A G T C T A G C G T A G T A C C G T A C T A G A T C G | Tbx20(T-box)/Heart-Tbx20-ChIP-Seq(GSE29636)/Homer | 1e-2 | -5.103e+00 | 0.0246 | 40.0 | 3.71% | 1131.6 | 2.42% | motif file (matrix) | svg |
| 107 | T G A C A G T C C G T A A C T G G T A C A C G T A C T G A C G T G A C T G A T C | Twist2(bHLH)/Myoblast-Twist2.Ty1-ChIP-Seq(GSE127998)/Homer | 1e-2 | -5.073e+00 | 0.0250 | 241.0 | 22.38% | 9010.3 | 19.28% | motif file (matrix) | svg |
| 108 | A C T G T G A C G T A C C G T A A G T C T A C G A C G T A C T G G T C A A G T C | NPAS2(bHLH)/Liver-NPAS2-ChIP-Seq(GSE39860)/Homer | 1e-2 | -5.033e+00 | 0.0258 | 145.0 | 13.46% | 5136.4 | 10.99% | motif file (matrix) | svg |
| 109 | T C G A G C A T A C T G C T G A A G T C T C A G G A C T G T A C C G T A A G C T A G T C G A T C | c-Jun-CRE(bZIP)/K562-cJun-ChIP-Seq(GSE31477)/Homer | 1e-2 | -5.017e+00 | 0.0260 | 38.0 | 3.53% | 1067.5 | 2.28% | motif file (matrix) | svg |
| 110 | G A C T G C A T C T A G C G A T G A T C T C G A C A T G G A T C | Tgif1(Homeobox)/mES-Tgif1-ChIP-Seq(GSE55404)/Homer | 1e-2 | -5.005e+00 | 0.0261 | 392.0 | 36.40% | 15322.7 | 32.79% | motif file (matrix) | svg |
| 111 | C G A T T G C A T G C A G A T C C G T A A C T G T G A C G A C T C A T G A C T G | Tcf21(bHLH)/ArterySmoothMuscle-Tcf21-ChIP-Seq(GSE61369)/Homer | 1e-2 | -4.976e+00 | 0.0266 | 141.0 | 13.09% | 4986.0 | 10.67% | motif file (matrix) | svg |
| 112 | C A G T A C T G T C A G T G C A G C T A A T G C T C G A A T C G G T C A T G C A | ZNF189(Zf)/HEK293-ZNF189.GFP-ChIP-Seq(GSE58341)/Homer | 1e-2 | -4.974e+00 | 0.0266 | 142.0 | 13.18% | 5026.6 | 10.76% | motif file (matrix) | svg |
| 113 | A C G T T C G A T C G A A G T C G T C A T A C G A T G C A C G T A C T G A G C T | Myf5(bHLH)/GM-Myf5-ChIP-Seq(GSE24852)/Homer | 1e-2 | -4.656e+00 | 0.0360 | 110.0 | 10.21% | 3811.7 | 8.16% | motif file (matrix) | svg |
| 114 | A T G C A G T C G C T A C G A T C G T A G C A T G C T A C G A T C T A G C A T G T G A C G T C A | CArG(MADS)/PUER-Srf-ChIP-Seq(Sullivan\_et\_al.)/Homer | 1e-2 | -4.620e+00 | 0.0370 | 35.0 | 3.25% | 990.1 | 2.12% | motif file (matrix) | svg |
