## Additional file S7-S10 for "Changes in microglia chromatin accessibility in aged female mice": Additional file S10_Homer_known motifs_loss.html

lostHM/ - Homer Known Motif Enrichment Results


### Homer Known Motif Enrichment Results (lostHM/)

Homer *de novo* Motif Results  
Gene Ontology Enrichment Results  
Known Motif Enrichment Results (txt file)  
Total Target Sequences = 193, Total Background Sequences = 47190

|  |  |  |  |  |  |  |  |  |  |  |  |
| --- | --- | --- | --- | --- | --- | --- | --- | --- | --- | --- | --- |
| Rank | Motif | Name | P-value | log P-pvalue | q-value (Benjamini) | # Target Sequences with Motif | % of Targets Sequences with Motif | # Background Sequences with Motif | % of Background Sequences with Motif | Motif File | SVG |
| 1 | T C G A T G A C G T A C C G T A C A G T T G A C A C G T A C T G A G C T A G C T | NeuroG2(bHLH)/Fibroblast-NeuroG2-ChIP-Seq(GSE75910)/Homer | 1e-28 | -6.614e+01 | 0.0000 | 98.0 | 50.78% | 7451.3 | 15.79% | motif file (matrix) | svg |
| 2 | T A C G T G C A A G T C C G T A A C G T T G A C A C G T A C T G A C T G G C A T | TCF4(bHLH)/SHSY5Y-TCF4-ChIP-Seq(GSE96915)/Homer | 1e-24 | -5.721e+01 | 0.0000 | 93.0 | 48.19% | 7515.4 | 15.92% | motif file (matrix) | svg |
| 3 | C T G A T G A C T G A C C G T A A C G T T G A C A G C T C T A G A C G T G A C T | Olig2(bHLH)/Neuron-Olig2-ChIP-Seq(GSE30882)/Homer | 1e-23 | -5.493e+01 | 0.0000 | 105.0 | 54.40% | 9823.2 | 20.81% | motif file (matrix) | svg |
| 4 | T C A G A C G T T C G A T A G C A G T C C G T A A C T G G T A C A C G T A C T G A T C G A G T C | Atoh1(bHLH)/Cerebellum-Atoh1-ChIP-Seq(GSE22111)/Homer | 1e-23 | -5.480e+01 | 0.0000 | 74.0 | 38.34% | 4901.2 | 10.38% | motif file (matrix) | svg |
| 5 | C T A G T C G A T G A C A G T C C G T A A C T G G T A C A C G T A C T G A C T G | BHLHA15(bHLH)/NIH3T3-BHLHB8.HA-ChIP-Seq(GSE119782)/Homer | 1e-21 | -4.935e+01 | 0.0000 | 82.0 | 42.49% | 6569.3 | 13.92% | motif file (matrix) | svg |
| 6 | T G A C A G T C C G T A A C T G G T A C A C G T A C T G A C G T G A C T G A T C | Twist2(bHLH)/Myoblast-Twist2.Ty1-ChIP-Seq(GSE127998)/Homer | 1e-20 | -4.758e+01 | 0.0000 | 94.0 | 48.70% | 8752.2 | 18.54% | motif file (matrix) | svg |
| 7 | T C A G T G A C G T A C C G T A A C G T T G A C A C G T T C A G A G C T G A C T | NeuroD1(bHLH)/Islet-NeuroD1-ChIP-Seq(GSE30298)/Homer | 1e-18 | -4.302e+01 | 0.0000 | 57.0 | 29.53% | 3591.4 | 7.61% | motif file (matrix) | svg |
| 8 | C T G A T C A G G T A C G C T A A C T G T G A C G C A T C A T G | SCL(bHLH)/HPC7-Scl-ChIP-Seq(GSE13511)/Homer | 1e-12 | -2.818e+01 | 0.0000 | 139.0 | 72.02% | 21941.3 | 46.49% | motif file (matrix) | svg |
| 9 | C T A G A G T C T A C G T A C G T G A C C G T A A C T G T A G C G C A T C A T G A T G C A G C T | Ascl1(bHLH)/NeuralTubes-Ascl1-ChIP-Seq(GSE55840)/Homer | 1e-8 | -2.023e+01 | 0.0000 | 60.0 | 31.09% | 6697.9 | 14.19% | motif file (matrix) | svg |
| 10 | C T A G A G C T G A C T C A T G A G T C A G T C G T C A C A G T C T A G T C A G G T A C C T G A T C G A G A T C T G A C | Rfx2(HTH)/LoVo-RFX2-ChIP-Seq(GSE49402)/Homer | 1e-8 | -1.869e+01 | 0.0000 | 12.0 | 6.22% | 307.6 | 0.65% | motif file (matrix) | svg |
| 11 | A C T G T C A G A G C T G A C T C A T G A G T C A G T C G C T A C G A T C T A G T C A G G T A C C T G A T C G A | Rfx1(HTH)/NPC-H3K4me1-ChIP-Seq(GSE16256)/Homer | 1e-7 | -1.812e+01 | 0.0000 | 17.0 | 8.81% | 735.4 | 1.56% | motif file (matrix) | svg |
| 12 | C A T G C T A G A G C T G A C T C A T G A G T C G A T C G C T A C G A T C T A G T C A G G T A C C T G A T C G A | X-box(HTH)/NPC-H3K4me1-ChIP-Seq(GSE16256)/Homer | 1e-7 | -1.699e+01 | 0.0000 | 13.0 | 6.74% | 436.0 | 0.92% | motif file (matrix) | svg |
| 13 | A G T C A T C G C T A G A G C T G A C T C T A G A G T C A G T C G C T A C A G T T C A G T C A G G A T C C T G A T C G A G A T C | RFX(HTH)/K562-RFX3-ChIP-Seq(SRA012198)/Homer | 1e-6 | -1.554e+01 | 0.0000 | 10.0 | 5.18% | 263.9 | 0.56% | motif file (matrix) | svg |
| 14 | T G A C C T G A C T A G C T G A C G T A A G T C C T G A A C G T G C A T T A G C G C A T A T C G G A C T G A C T G A T C | GRE(NR),IR3/RAW264.7-GRE-ChIP-Seq(Unpublished)/Homer | 1e-6 | -1.414e+01 | 0.0000 | 17.0 | 8.81% | 977.4 | 2.07% | motif file (matrix) | svg |
| 15 | T C G A A G T C C G T A A T C G T A G C A C G T A C T G A G C T A C G T A G T C | Ptf1a(bHLH)/Panc1-Ptf1a-ChIP-Seq(GSE47459)/Homer | 1e-5 | -1.255e+01 | 0.0001 | 79.0 | 40.93% | 12202.2 | 25.85% | motif file (matrix) | svg |
| 16 | C A G T T G C A A C G T A C T G C G T A A T C G C G A T T G A C C G T A A C G T | BATF(bZIP)/Th17-BATF-ChIP-Seq(GSE39756)/Homer | 1e-5 | -1.228e+01 | 0.0001 | 27.0 | 13.99% | 2508.3 | 5.31% | motif file (matrix) | svg |
| 17 | C A T G C T A G T C G A A C G T A C T G C G T A T A G C C G A T T G A C C G T A A G C T G A T C | Fra2(bZIP)/Striatum-Fra2-ChIP-Seq(GSE43429)/Homer | 1e-5 | -1.220e+01 | 0.0001 | 21.0 | 10.88% | 1651.1 | 3.50% | motif file (matrix) | svg |
| 18 | C G A T T G C A T G C A G A T C C G T A A C T G T G A C G A C T C A T G A C T G | Tcf21(bHLH)/ArterySmoothMuscle-Tcf21-ChIP-Seq(GSE61369)/Homer | 1e-5 | -1.209e+01 | 0.0001 | 36.0 | 18.65% | 3986.3 | 8.45% | motif file (matrix) | svg |
| 19 | T G A C G T A C C G T A A C T G T G A C C G A T A C T G A T C G A G C T T A C G T C G A T A G C G T A C C G T A A T C G T G A C G C A T A C T G A C T G A T G C | Twist(bHLH)/HMLE-TWIST1-ChIP-Seq(Chang\_et\_al)/Homer | 1e-4 | -1.122e+01 | 0.0003 | 11.0 | 5.70% | 526.4 | 1.12% | motif file (matrix) | svg |
| 20 | T C G A A C G T C A T G G C T A T A G C C G A T G T A C G C T A A C G T A T G C | AP-1(bZIP)/ThioMac-PU.1-ChIP-Seq(GSE21512)/Homer | 1e-4 | -1.086e+01 | 0.0004 | 28.0 | 14.51% | 2876.5 | 6.09% | motif file (matrix) | svg |
| 21 | A C T G C T A G T C G A C G A T C A T G G C T A A T C G C G A T G T A C G C T A A G C T G T A C | Fra1(bZIP)/BT549-Fra1-ChIP-Seq(GSE46166)/Homer | 1e-4 | -1.031e+01 | 0.0007 | 22.0 | 11.40% | 2025.1 | 4.29% | motif file (matrix) | svg |
| 22 | C T A G T C G A A C G T A C T G C G T A A T G C A C G T G T A C C G T A A G C T G A T C G T A C | Atf3(bZIP)/GBM-ATF3-ChIP-Seq(GSE33912)/Homer | 1e-4 | -1.028e+01 | 0.0007 | 25.0 | 12.95% | 2490.1 | 5.28% | motif file (matrix) | svg |
| 23 | C T A G T C G A G C A T C A T G G C T A T A G C C G A T G T A C C T G A A G C T | JunB(bZIP)/DendriticCells-Junb-ChIP-Seq(GSE36099)/Homer | 1e-4 | -9.344e+00 | 0.0016 | 21.0 | 10.88% | 2011.0 | 4.26% | motif file (matrix) | svg |
| 24 | G A C T C T A G A T G C A G T C G T C A T A C G A T G C A T C G | HIC1(Zf)/Treg-ZBTB29-ChIP-Seq(GSE99889)/Homer | 1e-3 | -8.762e+00 | 0.0028 | 58.0 | 30.05% | 8987.0 | 19.04% | motif file (matrix) | svg |
| 25 | T C A G G A C T C A G T C T G A A G C T C T A G G A C T T G C A C T G A A G T C | HLF(bZIP)/HSC-HLF.Flag-ChIP-Seq(GSE69817)/Homer | 1e-3 | -8.752e+00 | 0.0028 | 32.0 | 16.58% | 3951.9 | 8.37% | motif file (matrix) | svg |
| 26 | C G T A C T G A C T A G C G T A C G T A A G T C C G T A C A G T G C A T G T C A C G A T A C T G A C G T G C A T G A T C | PGR(NR)/EndoStromal-PGR-ChIP-Seq(GSE69539)/Homer | 1e-3 | -8.482e+00 | 0.0034 | 14.0 | 7.25% | 1104.4 | 2.34% | motif file (matrix) | svg |
| 27 | A G C T A C G T A C T G A T G C A G T C C G T A C T G A T A C G | NF1-halfsite(CTF)/LNCaP-NF1-ChIP-Seq(Unpublished)/Homer | 1e-3 | -8.268e+00 | 0.0041 | 52.0 | 26.94% | 7917.1 | 16.77% | motif file (matrix) | svg |
| 28 | T C G A A C G T A C G T C T G A G A T C T C A G G A C T G T C A C G T A A G C T G T C A C T A G A G C T A C G T T C G A | NFIL3(bZIP)/HepG2-NFIL3-ChIP-Seq(Encode)/Homer | 1e-3 | -7.719e+00 | 0.0068 | 24.0 | 12.44% | 2770.1 | 5.87% | motif file (matrix) | svg |
| 29 | A T G C A G T C G T A C A G C T T C G A C T A G G A T C C T G A G T C A A G T C G C T A T C A G | Rfx5(HTH)/GM12878-Rfx5-ChIP-Seq(GSE31477)/Homer | 1e-3 | -7.184e+00 | 0.0112 | 17.0 | 8.81% | 1716.3 | 3.64% | motif file (matrix) | svg |
| 30 | A G C T C A T G G C A T G A T C T G C A C T A G G A T C A C G T | Tgif2(Homeobox)/mES-Tgif2-ChIP-Seq(GSE55404)/Homer | 1e-3 | -7.116e+00 | 0.0116 | 85.0 | 44.04% | 15530.2 | 32.90% | motif file (matrix) | svg |
| 31 | T G A C A T G C C G T A A T C G A T G C C A G T C A T G A C T G A G T C G T A C | HEB(bHLH)/mES-Heb-ChIP-Seq(GSE53233)/Homer | 1e-3 | -6.921e+00 | 0.0136 | 50.0 | 25.91% | 7960.2 | 16.87% | motif file (matrix) | svg |
| 32 | C T A G T A C G G A C T T G C A T G C A C G A T T A C G C T G A T C G A C T G A | Hoxa10(Homeobox)/ChickenMSG-Hoxa10.Flag-ChIP-Seq(GSE86088)/Homer | 1e-2 | -6.667e+00 | 0.0170 | 25.0 | 12.95% | 3168.8 | 6.71% | motif file (matrix) | svg |
| 33 | A G T C G A T C A G T C C G T A A T C G C A G T A G T C G T A C C T G A A C T G T C A G A G C T A G C T A G C T A G C T | PRDM15(Zf)/ESC-Prdm15-ChIP-Seq(GSE73694)/Homer | 1e-2 | -6.416e+00 | 0.0212 | 33.0 | 17.10% | 4723.2 | 10.01% | motif file (matrix) | svg |
| 34 | C T A G T C G A C G A T A C T G C G T A T A C G A G C T T G A C G C T A A C G T G A T C T A G C | Fosl2(bZIP)/3T3L1-Fosl2-ChIP-Seq(GSE56872)/Homer | 1e-2 | -6.338e+00 | 0.0222 | 12.0 | 6.22% | 1078.2 | 2.28% | motif file (matrix) | svg |
| 35 | C A G T T C A G G A T C A C T G A C G T C T A G A C T G A C T G G A C T C T A G | Egr1(Zf)/K562-Egr1-ChIP-Seq(GSE32465)/Homer | 1e-2 | -6.263e+00 | 0.0233 | 17.0 | 8.81% | 1871.7 | 3.97% | motif file (matrix) | svg |
| 36 | T A G C G T A C C G T A C T A G A C T G T G C A C G T A A T G C C G T A A T C G | AR-halfsite(NR)/LNCaP-AR-ChIP-Seq(GSE27824)/Homer | 1e-2 | -6.094e+00 | 0.0268 | 96.0 | 49.74% | 18594.1 | 39.40% | motif file (matrix) | svg |
| 37 | T G C A A G C T A C G T C T A G G A T C C T A G G A T C G T C A C T G A A G T C | CEBP(bZIP)/ThioMac-CEBPb-ChIP-Seq(GSE21512)/Homer | 1e-2 | -5.454e+00 | 0.0495 | 23.0 | 11.92% | 3100.3 | 6.57% | motif file (matrix) | svg |
| 38 | C G T A A C G T A C G T A C G T A C G T A G T C A G T C C T G A A G C T A G C T | NFAT(RHD)/Jurkat-NFATC1-ChIP-Seq(Jolma\_et\_al.)/Homer | 1e-2 | -5.435e+00 | 0.0495 | 32.0 | 16.58% | 4829.9 | 10.23% | motif file (matrix) | svg |
| 39 | C T A G T C G A A C G T A C T G C G T A T A G C C G A T G T A C C G T A A G C T G A T C G T A C | Jun-AP1(bZIP)/K562-cJun-ChIP-Seq(GSE31477)/Homer | 1e-2 | -5.380e+00 | 0.0506 | 9.0 | 4.66% | 766.8 | 1.62% | motif file (matrix) | svg |
| 40 | T G A C C T G A C T A G T C G A C T G A A T G C C G T A A C T G G C A T G T A C G C A T A T C G G C A T A G C T G A T C | PR(NR)/T47D-PR-ChIP-Seq(GSE31130)/Homer | 1e-2 | -5.292e+00 | 0.0538 | 64.0 | 33.16% | 11654.1 | 24.69% | motif file (matrix) | svg |
| 41 | C G A T G A C T C G A T T C A G G A C T A C G T C A G T C T G A G A C T G A C T A G C T C G A T A C T G A T C G G T A C G C T A | NF1:FOXA1(CTF,Forkhead)/LNCAP-FOXA1-ChIP-Seq(GSE27824)/Homer | 1e-2 | -5.155e+00 | 0.0602 | 5.0 | 2.59% | 275.2 | 0.58% | motif file (matrix) | svg |
| 42 | C T G A A G C T A C G T A C G T A G T C G A C T G A C T C T G A C T G A C T A G C G T A C G T A | STAT6(Stat)/CD4-Stat6-ChIP-Seq(GSE22104)/Homer | 1e-2 | -4.930e+00 | 0.0736 | 21.0 | 10.88% | 2867.7 | 6.08% | motif file (matrix) | svg |
| 43 | C T G A T C A G A G T C C G T A A T C G A T G C C G A T A C T G A G T C G A C T A T C G A G T C | MyoD(bHLH)/Myotube-MyoD-ChIP-Seq(GSE21614)/Homer | 1e-2 | -4.759e+00 | 0.0853 | 22.0 | 11.40% | 3101.2 | 6.57% | motif file (matrix) | svg |
| 44 | T G C A A G C T C T G A A T C G G A C T C T A G G T A C G A T C G T C A A G T C G T A C G A C T C T A G A T C G G C A T C A T G C A T G G A T C G T A C C T G A | CTCF(Zf)/CD4+-CTCF-ChIP-Seq(Barski\_et\_al.)/Homer | 1e-2 | -4.716e+00 | 0.0871 | 7.0 | 3.63% | 563.1 | 1.19% | motif file (matrix) | svg |
| 45 | C A T G A G T C C T G A T G A C A T C G G A C T G T C A A G T C T A G C G A T C | HIF2a(bHLH)/785\_O-HIF2a-ChIP-Seq(GSE34871)/Homer | 1e-2 | -4.623e+00 | 0.0935 | 11.0 | 5.70% | 1180.8 | 2.50% | motif file (matrix) | svg |
| 46 | T C A G A G C T A C G T A C G T G T A C G A T C C G T A C T A G C A T G G T C A C G T A T C G A | STAT4(Stat)/CD4-Stat4-ChIP-Seq(GSE22104)/Homer | 1e-2 | -4.622e+00 | 0.0935 | 30.0 | 15.54% | 4707.4 | 9.97% | motif file (matrix) | svg |
